# Systemic inflammation recruits fast-acting anti-inflammatory innate myeloid progenitors from BM into lymphatics

**DOI:** 10.1101/2021.01.20.427403

**Authors:** Juana Serrano-Lopez, Shailaja Hegde, Sachin Kumar, Josefina Serrano, Jing Fang, Ashley M. Wellendorf, Paul A. Roche, Yamileth Rangel, Léolène J. Carrington, Hartmut Geiger, H. Leighton Grimes, Sanjiv Luther, Ivan Maillard, Joaquin Sanchez-Garcia, Daniel T. Starczynowski, Jose A. Cancelas

## Abstract

Innate immune cellular effectors are actively consumed during systemic inflammation but the systemic traffic and the mechanisms that support their replenishment remain unknown. Here we demonstrate that acute systemic inflammation induces the emergent activation of a previously unrecognized system of rapid migration of granulocyte-macrophage progenitors and committed macrophage-dendritic progenitors, but not other progenitors or stem cells, from bone marrow (BM) to lymphatic capillaries. The progenitor traffic to the systemic lymphatic circulation is mediated by Ccl19/Ccr7 and is NFκB independent, Traf6/IκB-kinase/SNAP23 activation which is responsible for the secretion of pre-stored Ccl19 by a subpopulation of CD205^+^/CD172a^+^ conventional dendritic cells type 2 (cDC2) and upregulation of BM myeloid progenitor Ccr7 signaling. The consequence of this progenitor traffic is anti-inflammatory with promotion of early survival and initiation of replenishment of lymph node cDC.

## INTRODUCTION

Bacterial infections represent one of the major threats for the human immune system and can lead to sepsis or death (Martin et al., 2003). A functional immune response is a key factor to control the outcome of bacterial infections. Therefore, the human immune system has evolved several effector mechanisms to fight bacterial infections which involve the innate and the adaptive arms of the immune system. Antigen presentation is an essential mechanism of activation that requires crosstalk between the innate and adaptive immune system to fight bacterial infections. Dendritic cells are short-lived professional antigen presenting cells (APC) and their life span is further reduced during the inflammatory response to pathogens (Kamath et al., 2002). Upon inflammation, primed APC thus need to be replaced.

During inflammation, systemic signals alert and activate bone marrow (BM) myeloid differentiation (Baldridge et al., 2010; Essers et al., 2009; Nagai et al., 2006). Inflamed secondary lymphoid organs such as lymph nodes (LN) recruit antigen-presenting dendritic cells (DC) (Legler et al., 1998; Luther et al., 2000; Saeki et al., 1999), while pathogen-associated molecular pattern signals (PAMPs) trigger migration of tissue-resident DC to the LN (Kaisho and Akira, 2001; Sallusto and Lanzavecchia, 2000). Circulation of hematopoietic stem cells and progenitors (HSC/P) that enter the lymphatic vessels from the peripheral blood (PB) with ability to amplify APCs has been described (Massberg et al., 2007). However, the circuits used by these HSC/P populations, their characterization and the cellular and molecular mechanisms that regulate this traffic in inflammatory conditions have not been addressed in detail.

Lymphatics form part of an open circulatory system that drains cells and interstitial fluid from tissues. Recently, bone lymphatic endothelial cells have been shown to arise rapidly from pre-existing regional lymphatics in inducible bone-expressing *Vegfc* transgenic mice through Vegfr3, osteoclast activation and bone loss (Hominick et al., 2018; Monroy et al., 2020). Acute endotoxemia is associated with osteoclast activation and bone loss (Hardy and Cooper, 2009; Nason et al., 2009). We postulated the pre-existence of an anatomical and functional patent circuit that communicates BM and lymphatic tissues that can be induced upon severe inflammatory conditions like endotoxemia.

Our work identifies an emergent traffic of DC-biased myeloid progenitors through direct transit from BM to bone lymphatic capillaries. This traffic is highly activated in endotoxic inflammation. In human reactive lymphadenitis or just after a single immune endotoxic challenge, such as following lipopolysaccharide (LPS) stimulation in mice, a massive mobilization of myeloid progenitors from the BM to lymph and retention in the LN takes place. The mobilization is rapid, prior to their appearance in PB. LPS simultaneously induces cell-autonomous Ccr7 expression on granulo-macrophage progenitors (GMP) and macrophage-dendritic progenitors (MDP), and a non-cell autonomous myeloid cell-dependent secretion of Ccl19 in the LN. In vivo blockade of LPS signaling in mature myeloid cells, deletion of hematopoietic Ccl19 or neutralization of Ccr7 completely abrogated the GMP/MDP migration from the BM to the LN and increased acute inflammation associated mortality. Moreover, genetic and pharmacological approaches revealed that Traf6-Irak1/4-Ubc13-IκB kinase (IKK) signaling mediates NF-κB-independent-SNAP23 phosphorylation and secretion of pre-formed Ccl19 from a specific population of conventional dendritic cells (cDC). These findings indicate that inflammation results in mobilization of cDC-forming cells directly from the BM to the lymph and LN. As such, emergent myeloid lineage mobilization from the BM to lymph may be important in inflammation by acutely replenishing antigen-presenting cells in lymph tissues and impairing the inflammatory signaling responsible for mortality in endotoxemia.

## RESULTS

### Inflammation associates with emergent migration of myeloid progenitors, but not HSC, from BM to lymphatics

To determine whether there is a circulation of HSC/P to human LN, we prospectively analyzed the presence of side population (SP) cells in LN biopsies (**Figure S1A**) obtained from lymphadenitis and lymphoma patients at diagnosis. Human and murine SP cells, with ability to extrude the dye Hoechst 33342 through upregulated activity of multidrug resistance protein complexes (Zhou et al., 2001) in BM and other tissues (Brusnahan et al., 2010; Challen and Little, 2006; Goodell et al., 1996) are enriched in long-term reconstituting HSC and other more committed populations of progenitors (Matsuzaki et al., 2004; Weksberg et al., 2008). We found a SP population at a frequency higher than 0.01% in 36 out of 64 LN biopsies (53.12%). However, the content of SP cells in the LN did correlate with the LN histological diagnosis. The elevated frequency of SP cells in LN did correlate with the LN histological diagnosis (**Figure 1A**) but not to the anatomical location of the lymphadenopathy (**Figure S1B; Table S1**). The accumulation of SP cells was significantly higher in LN from lymphadenitis patients than in lymphoma patients. Further dissection based on histological classifications by independent pathology analysis resulted in the lymphadenitis specimens being sorted into distinct histological categories which corresponded to follicular lymphadenitis with paracortical predominance (FL), granulomatous lymphadenitis (GL), and lymphadenopathies with histological or molecular evidence of viral etiology (viral lymphadenitis, VL). Interestingly, FL and GL LN contained a median of 0.2% SP cells with a range from <0.01% to ∼40%, which was significantly higher than the content of SP cells in VL, Hodgkin’s lymphoma, and non-Hodgkin’s lymphoma LN (**Figure 1A; Figure S1A, Table S1**). The existence of myeloid-committed hematopoietic progenitors was confirmed in myeloid colony-forming cell unit (CFU) assays (**Figure S1C**) performed on samples from patients with FL. These data show that non-viral inflammatory lymphadenitis results in a significantly increased frequency of primitive hematopoietic cells in LN, while it does not reveal the type of progenitor cells. To confirm whether LN SP cells indeed contained HSC/P, we first sorted LN SP cells from patients with reactive lymphadenitis and plated them in semisolid cultures containing rhIL-3, rhIL-6 and rhSCF cytokines CFU analysis demonstrated that SP cells were indeed capable of producing myeloid colonies (**Figures S1D**), while non-SP cells were devoid of measurable CFU-forming ability (data not shown). Immunophenotypic analysis of SP-derived progenitors was also consistent with enrichment of a heterocellular population of CD34 and CD133 expressing granulocyte-, granulocyte-macrophage-, and cDC-biased progenitors (Bornhauser et al., 2005; Gorgens et al., 2013) (**Figure S1D**). The vast majority of CD45^+^/CD34^+^ cells co-expressed CD133^+^, and the CD45^+^/CD34^−^ population was split ∼50:50 into CD133^+^ and CD133^−^ cells (**FigureS1D**). In combination, these data show the accumulation of a myeloid-committed HSC/P population in human lymphadenitis. Adult inflammatory LN tissues therefore contain an increased number of myeloid-committed HSC/P. This increase can result from either the recruitment of these cells to LN via the bloodstream or the expansion of otherwise rare and already resident myeloid-committed HSC/P in these LN.

**Figure 1.**
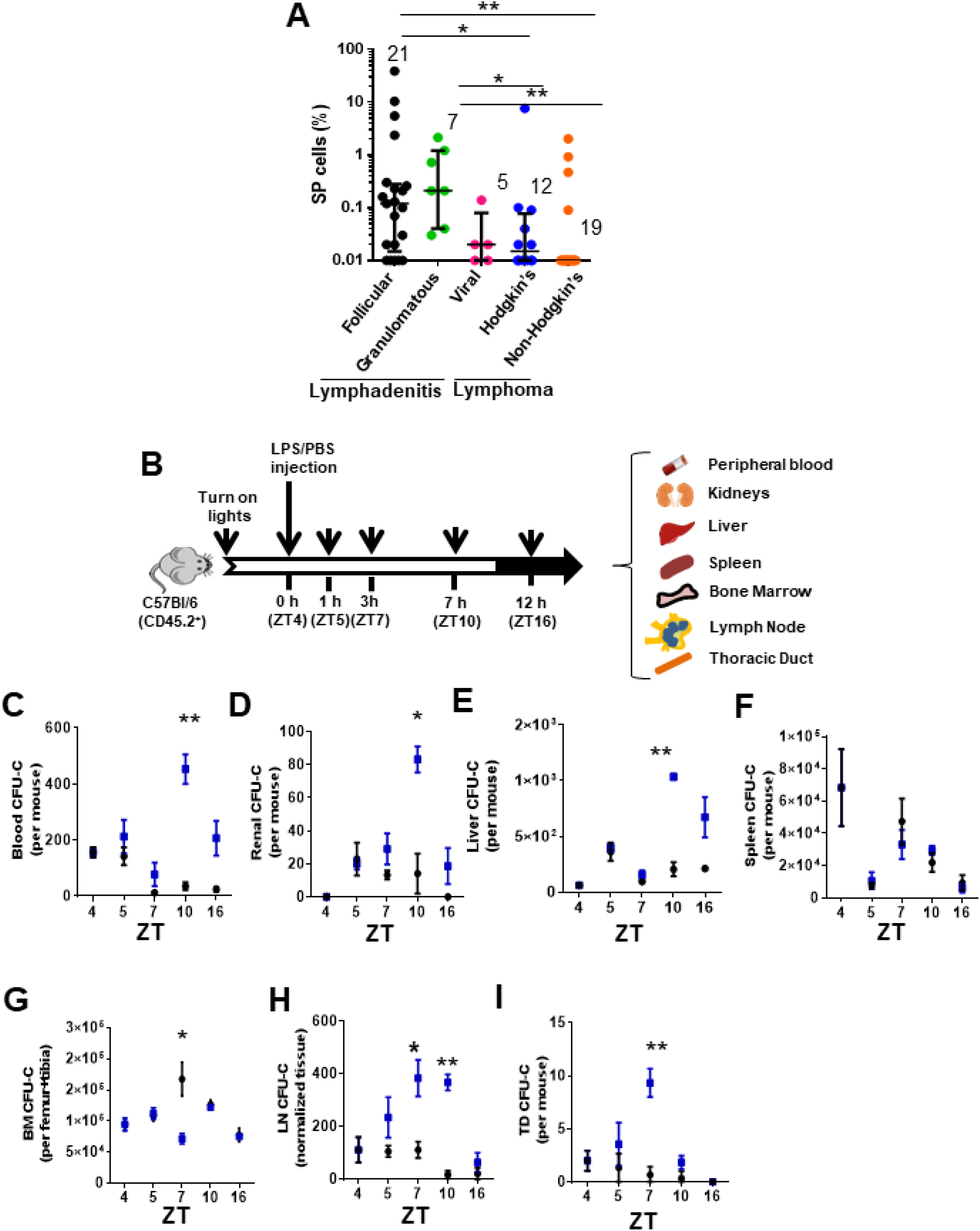
Inflammation induces early mobilization of HSC/Ps to lymph organs in humans and mice. (A) Content of SP cells in human LN by flow cytometry. LN biopsies had been blindly identified histologically as lymphadenitis, subcategorized in follicular (FL, black circles n=21), granulomatous (GL, green circles N=7) and viral (VL, pink circles n=5) and lymphomas, subcategorized in Hodgkin’s lymphoma (HL, blue circles n=12) or non-Hodgkin’s lymphoma (NHL, orange circles n=19). (B) Strategy for LPS administration and collection of tissues (blood, kidneys, liver, spleen, BM, LN and TD) at specific times. LPS or vehicle control PBS was administered at the early rest phase into C57Bl/6 (CD45.2^+^) mice and tissue specimens were collected before (ZT4), 1h (ZT5), 3 h (ZT7), 6 h (ZT10) or 12h (ZT16) later. (C-I). Myeloid colony-forming-cell unit (CFU-C) content in the collected tissues at different circadian cycle times. CFU-C contained in blood (C), kidneys (D) liver (E), spleen (F), BM (G), LN (H) and TD (I) in response to PBS (black circles) or LPS (blue squares) at different circadian cycle times (n=3-4 mice per time point and treatment). Results are shown as mean ± SEM. *P< 0.05, **P<0.01.

The release of HSC/P from BM into the bloodstream follows circadian cycles (Mendez-Ferrer et al., 2008) controlled by the activity and fate of inflammatory cells (Casanova-Acebes et al., 2013; Chang et al., 2014). We postulated that if inflammation is responsible for the recruitment of HSC/P to the LN and possibly other organs, we should be able to recapitulate the process of BM egression, migration, and organ retention, in an inflammatory murine model wherein the HSC/P migration process is highly conserved. Since the largest content of HSC/P in human LN was found in biopsies from patients with lymphadenitis, we generated a mouse model of Gram-negative sepsis by injection of *E. coli* LPS into C57Bl/6 mice at the early timepoint of the circadian HSC/P mobilization cycle (*zeitgeber* time, ZT) (Bellet et al., 2013) (**Figure1B**). *E. coli* LPS is able to activate a large number of Toll-like receptors (TLR), which result in high-level activation of the inflammatory signaling cascade (Beutler and Rietschel, 2003). LPS is also a well-known inducer of HSC/P mobilization to PB(Cline and Golde, 1977; Velders et al., 2004; Vos et al., 1972; Vos and Wilschut, 1979). In our experiments, the circadian mobilization pattern of HSC/P in the PB was severely modified by the administration of LPS, with the increase in HSC/P appearing later and peaking at ZT10, 6 hours post-administration (**Figure1C**), coincident with an increased neutrophil count in the PB (**Figure S1E**). We observed similar kinetics of an increased numbers of HSC/P in the highly vascularized kidney and liver tissues after LPS administration (**Figure 1D-E**), suggesting that the presence of HSC/P in these tissues closely paralleled their presence in the PB. Interestingly, LPS did not elicit a significant change in the level of splenic HSC/P within the first 12 hours after inflammation (Esplin et al., 2011; Wright et al., 2001) (**Figure 1F**).

Notably, when the HSC/P content was reduced in the BM, the kinetics of their subsequent mobilization to the PB was discordant. The BM HSC/P content decreased, which supports the migratory nature of the increased HSC/P in the PB (**Figure 1G**); yet the nadir of the BM HSC/P content occurred as early as 3 hours after LPS administration (at ZT7), returning to normal values by 6 hours (ZT10, **Figure 1G**). The time lapse between the loss of retention of HSC/P in the BM and their presence in the PB circulation suggested that the migration of HSC/P from the BM to the PB required an intermediate step of circulation through other tissues. Based on an earlier description of a lymphatic circulation of HSC/P (Massberg et al., 2007), we hypothesized that this delay in the appearance of HSC/P in the PB was due to an intermediate transit of HSC/P through the lymphatic circulation. Indeed, the lymphatic circulation in LPS-treated animals did show a significant increase in the levels of circulating HSC/P in the LN and the thoracic duct compared to controls that closely mirrored the decline of HSC/P in the BM (**Figures 1H-I**).

We next characterized the type of primitive cell populations migrating into the LN via the lymphatic circulation. We first analyzed whether the content of immunophenotypically identifiable BM HSC populations changed concomitantly with the progenitor population changes previously described. LPS induced expansion of BM Lin^−^/c-kit^+^/Sca-1^+^ (LSK) and immunophenotypically identified long-term (LT)-HSC, short-term (ST)-HSC and multipotential progenitors (MPP) populations at later time points (Z10-ZT16) (**Figures S1F and S1H-K**) with no changes in the BM HSC content by ZT7, suggesting a differential effect of LPS signaling on the HSC population. Interestingly, the reduction in the BM content of progenitors was not homogenous throughout the hematopoietic progenitor populations. Confirming the egress of BM CFU-GM described above, the GMP population was significantly decreased by ZT7 (**Figures S1G, L**), while the content of immunophenotypically defined common myeloid progenitors (CMP) only declined by ZT10 (**Figures S1G, M**), and the megakaryocyte-erythroid progenitor (MEP) content was increased (**Figures S1G, N**), resulting in no significant net changes in the content of BM Lin^−^/c-kit^+^/Sca1^−^ (LK) cells (**Figures S1G, O**).

Functional in vivo assays of LN cell suspensions obtained at ZT7 demonstrated that the accumulation of progenitors in LN did not contain any significant numbers of long-term or medium-term repopulating HSC. The analysis of competitive-repopulating units (CRU) in the LN (**Figure S2A**) demonstrated that inflamed LN did not contain increased levels of repopulating cells by ZT7 (**Figure S2B**). LN contained a transient, ST-myelopoietic progenitor population without medium- or long-term multilineage repopulation ability (**Figure S2C**). Lineage analysis of donor-derived circulating cells demonstrated no significant change in T-cell transfer (**Figure S2D**), and a diminished transfer of B-cells into the lethally irradiated recipients (**Figure S2E**) indicating the presence of adoptively transferred lymphoid cells and the absence of mobilization of competent lymphoid progenitors to the inflamed LN. Furthermore, LN SP cells from LPS-treated mice are enriched in LK cells and depleted in LSK cells (**Figure S2F**). LN SP cells contain exclusively ST-repopulating progenitors with the ability to differentiate into myeloid cells (data not shown) and are depleted from any significant 8-16 weeks engrafting HSC, as assessed using CRU assays (**Figures S2F-G**), unlike their BM SP counterparts which are enriched in LSK cells and LT-repopulating activity (**Figures S2F, H**). These results confirmed that, similar to human inflammatory LN, the LN SP cells from mice treated with *E. Coli* LPS accumulate in inflamed LN, contain progenitors, and are depleted of stem cell activity. Altogether, these data indicate that LPS induces a selective lymphatic circulation of myeloid committed progenitors, but not other types of HSC/P populations.

To explore the nature of the circuit of the myeloid progenitor migration to LN, we first analyzed the ability of HSC/P to seed LN in non-myeloablated mice. For this experiment, we labeled C57Bl/6-BM-derived lineage negative (Lin^−^) cells containing the HSC/P fraction with the lipophilic dye 1,1’-dioctadecyl-3,3,3’,3’-tetramethylindocarbocyanine perchlorate (DiI), and adoptively transferred them into un-manipulated lymphatic vessel reporter Lyve1-eGFP mice. Lyve1-GFP knock-in mice display enhanced green fluorescent protein (eGFP) fluorescence driven by the promoter/enhancer of the lymphatic vessel hyaluronan receptor 1 Lyve-1 identifying lymphatic endothelial cells (Pham et al., 2010). We analyzed the 17-hour homing of Lin^−^ cells to the BM and LN (**Figure 2A)**. The homing of cells to the BM in mice treated with LPS was reduced by ∼65% compared with their vehicle-treated counterparts (**Figure 2B**). To determine whether Lin^−^/DiI^+^ homed cells leave BM in response to LPS, we first determined the existence of transcortical vessels (**Figure 2C**)(Gruneboom et al., 2019) and the presence of lymphatic vasculature inside the bone by two-photon microscopy in mice either treated with PBS or LPS (**Figures 2D-E and Movies S1-2).** Lyve1-GFP knock-in mice revealed that the bone of LPS-treated animals contains a Lyve1+ network, which was only rarely identified in PBS-treated mice (**Figure 2D**), suggesting that LPS induced inflammation may upregulate the expression of the hyaluronan receptor Lyve1 and render patent a pre-existing network of Lyve1+ bone cells. Interestingly, Lin^−^/DiI^+^ homed cells were located closer to the endosteum in response to LPS at as early as 1.5 hours after administration of LPS (**Figure 2E)**. Quantification of the distance of Lin^−^/DiI^+^ homed cells to endosteum area showed significant differences between PBS and LPS treatment, indicating increased proximity to the endosteum area after LPS (**Figure 2F)**. We found, albeit at a very low frequency, tiny lymphatics scattered and projected inside the bone (**Figure 2G and Movie S3**). On the other hand, the seeding of BM-derived Lin^−^/DiI^+^ cells into LN increased ∼3-fold which mirrored the decline in BM homing (**Figure 2H-J**). Histological analysis of BM-derived Lin^−^/DiI^+^ cells within the LN by confocal microscopy showed that the migrated HSP/C are spatially positioned in the cortex area surrounding primary follicles (**Figure 2I)**, consistent with localization in T-cell zone for antigen presentation. These findings strongly suggest that the rapid egress of hematopoietic progenitors from BM during inflammation may indeed occur through bone lymphatics draining into LN.

**Figure 2.**
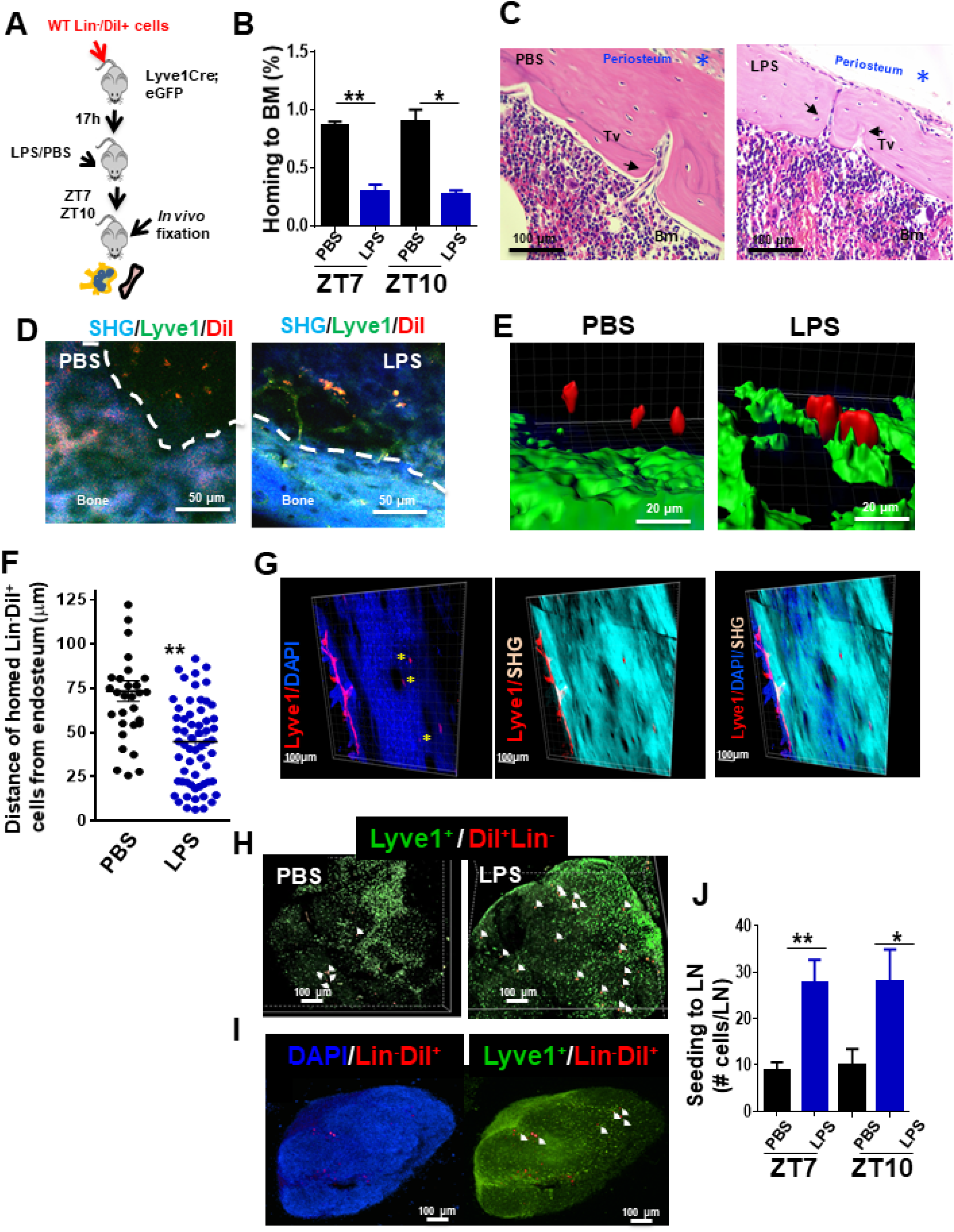
Draining of BM-derived lineage negative cells into lymphaticstem. (A) Schema for adoptive transfer of BM-derived Lineage negative cells (Lin^−^) labeled with 1,1’-dioctadecyl-3,3,3’,3’-tetramethylindocarbocyanine perchlorate (DiI) dye into lymphatic endothelium reporter Lyve1-eGFP^+^ mice. By 16 hours after cell transplantation, LPS or PBS were administered into the Lyve1-eGFP^+^ mice at ZT4 (0h). BM and LN cells were analyzed at ZT5.5 (1.5h), ZT7 (3h) and ZT10 (6h) for labeled Lin^−^ cells (Lin^−^/DiI^+^). (B) Frequency of Lin^−^/DiI^+^ homed to BM (solid bars) (mosaic bars) after PBS (black solid bar) or LPS (blue solid bar) administration at ZT7 (3h) and ZT10 (6h). N=3 mice per time point and group in two independent experiments. Graph represent mean ± SEM. *P<0.05, **P<0.01. (C) Representative 3D reconstruction images of the whole bone by two-photon microscopy showing lymphatic vessels from Lyve1 surface marker (red), nuclei with DAPI (Blue) and cortical bone with second harmonic signal (SHG, light blue). The scale bars, 100 μm. (D, i-iv) Intravital two-photon microscopy imaging (IVM) of long bones from Lyve1-eGFP mice showing lymphatic vessels (displayed in green) near to the surface of the bone (blue, detected by SHG signal) and homed Lin^−^/DiI^+^ cells (displayed in red). (i,iii) IVM of PBS specimen. (ii,iv) IVM of LPS specimen. (E,F) Analysis and quantification of the distance of homed Lin^−^/DiI^+^ (red) cells to Lyve1-EGFP+ (green) cells after PBS/LPS administration at ZT5.5 (1.5h) analyzed by Imaris 7.7.2 software. (G) Two-photon microscopy examples of images of longitudinal femoral sections stained with anti-Lyve1 antibody and DAPI, and analyzed for specific fluorescence signal and SHG for cortical bone. (H-I) Representative of 3D reconstitution images of PBS- and LPS-treated LN tissues (H) and cross-sections of LPS-treated LN (I) analyzed by confocal microscopy showing the location of mobilized Lin^−^/DiI^+^ cells (red; nucleus stained by DAPI in blue) in relation with Lyve-1+ cells (green; nucleus stained by DAPI in blue). The Z-stack dimensions of upper panels were: X=1266.95μm, Y=1266.95μm and Z=344μm. Calibrate: XY=2.47μm and Z=4μm. Resolution: 512 x 512 x 86. The Z-stack dimensions of lower panels were: X=1259.36μm, Y=1259.36μm and Z=132μm. Calibrate: XY=2.46μm and Z=4μm. Resolution of images were: 512 x 512 x 86. (J) Absolute count of mobilized Lin^−^/DiI^+^ cells counted within LN at ZT7 (3h, solid bars) and ZT10 (6h, mosaic bars) after PBS/LPS administration. N=4-14 lymph nodes analyzed per time point and graph represent mean ± SEM. *P< 0.05, **P<0.01.

### Systemic inflammation recruits dendritic cell-committed phenotypic progenitors to LN

To determine the potential of the myeloid progenitors mobilized to the LN, we further determined their in vitro and in vivo differentiation profile. To this end we analyzed the differentiation capabilities of LN myeloid progenitors in specific-cytokine driven clonal assays in methylcellulose assays. The majority of the differentially accumulated myeloid progenitors in LN by 3 hours post-administration of LPS were granulocyte-macrophage progenitors (CFU-GM) and in a much lesser degree unipotent granulocyte progenitors (GPs, CFU-G) with no differential accumulation of unipotent macrophage progenitors (MPs, CFU-M) (**Figure 3A**). Next, we investigated whether GMP were able to home and migrate to LN after in vivo administration of LPS. For this purpose, we adoptively transferred sorted β-actin/eGFP transgenic GMPs into congenic mice. Transgenic GMPs were allowed to home to the BM and after 17 hours recipient mice were treated with a single dose of LPS or vehicle control. On day 7 after PBS or LPS administration, murine BM and LN were analyzed for donor-derived granulocytes (Gr1^++^/CD11b^+^/CD11c^neg^), macrophages (Gr1^dim^/CD11b^+^/CD11c^neg^) and cDC (Gr1^neg^/CD11b^+^/CD11c^+^) by flow cytometry (**Figure 3B**). We found that LPS induced differential donor-derived specific GMP differentiation towards the formation and retention of cDC in LN (**Figure 3C**), but not in the BM (**Figure 3D**). The content of macrophages and granulocytes did not significantly change with LPS in either LN or BM (**Figures 3C-D**) confirming the specific nature of the cDC differentiation of mobilized GMP in LPS-treated mice, similar to our observations in human lymphadenitis.

**Figure 3.**
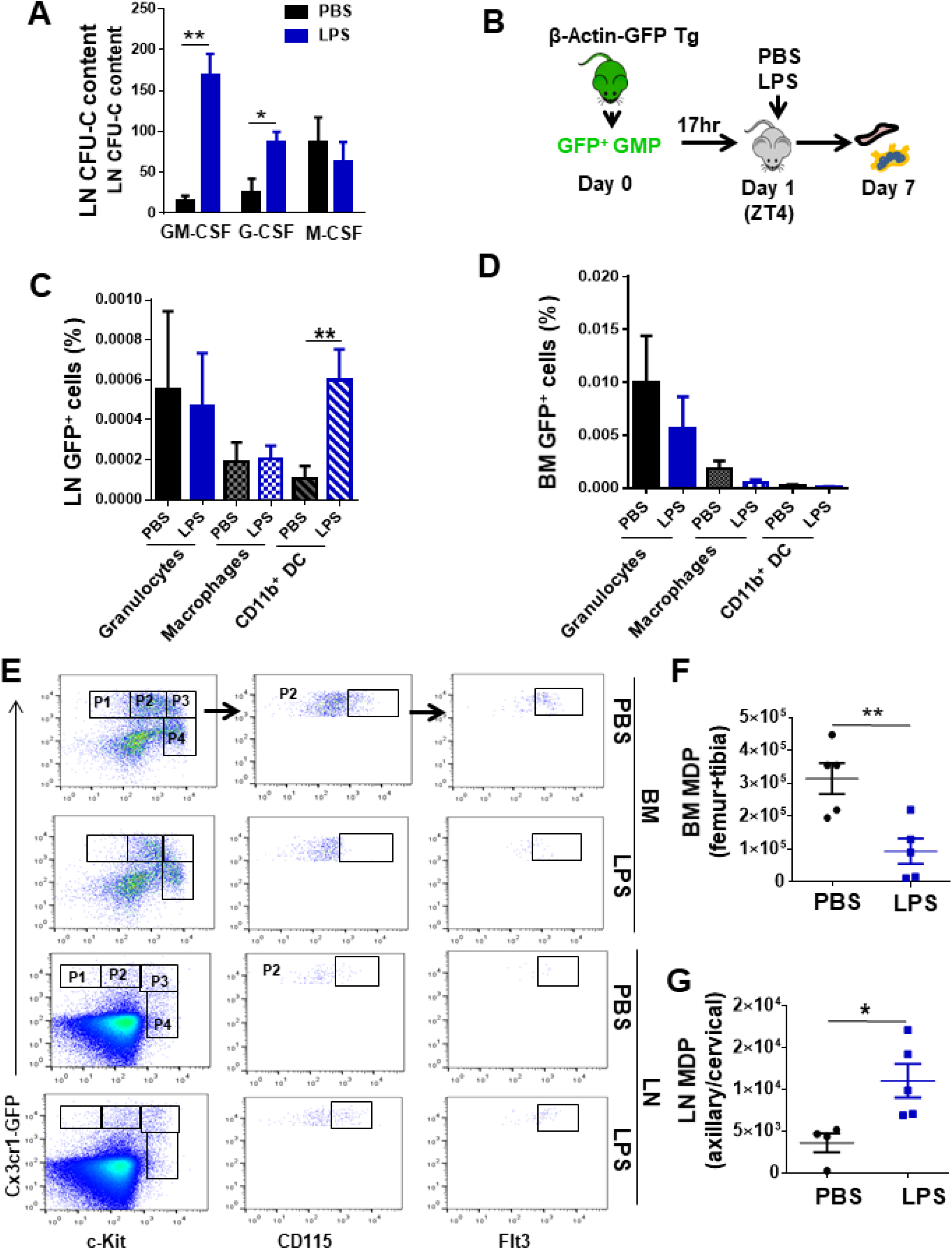
LN-mobilized GMP preferentially differentiate into dendritic cells. (A) Comparative quantification of the content of bipotent and unipotent myeloid progenitors in LN at ZT7 (3h) after PBS/LPS administration at ZT4 (0h). (B) Schema of isolation and transfer of BM-derived GMP from β-Actin-GFP reporter mice (CD45.2^+^) into C57Bl/6 (CD45.2^+^) mice (2×10^5^ GFP^+^-GMP cells/mouse, n=3 mice per group). After BM homing (17h), mice were treated with single and low dose of LPS (5mg/Kg) at ZT4 (day1) and 7 days later BM and LN tissues were analyzed for GFP expression in myeloid populations by flow cytometry. (C-D) Graphs represent the percentage of GFP^+^-GMP differentiated to granulocytes (solid bars, Gr1^++^CD11b^+^CD11c^−^), macrophages (left mosaic bars, Gr1^dim^CD11b^+^CD11c^neg^) and cDC (right mosaic bars, Gr1^−^CD11b^+^ CD11c^+^) 7 days post-transferring after PBS/LPS administration by flow cytometry into LN (C) and BM (D). Values represent as mean ± SEM. **P<0.01. (E) FACS strategy for MDP content in BM and LN tissues from Cx3cr1^gfp+^ reporter mice (4-5 mice per group). Phenotypically, MDP are defined as lineage-negative with high expression of the chemokine receptor Cx3cr1, c-fms (CD115) and Flt3 (P2), and intermediate expression of c-Kit. (F-G) Graphs show absolute numbers of MDP present in BM (F) and LN (G) 3 hour later (ZT7 [3h]) after PBS (black circles) or LPS (blue squares) administration. Values represent mean ± SEM *P<0.05, **P<0.01.

To elucidate whether the LN cDC content was dependent on migration of committed cDC precursors opposed to local specification of migrated macrophages or macrophage progenitors, we analyzed the migration of macrophage-dendritic precursors (MDP). BM MDP is a progenitor population that can differentiate into monocytes/macrophages or directly into cDCs without intermediate macrophage specification (Fogg et al., 2006; Geissmann et al., 2003; Waskow et al., 2008). MDP are characterized by high expression of the chemokine receptor Cx3cr1, c-fms (CD115) and Flt3, and intermediate expression of c-Kit. Serial gating of Lineage^−^/Cx3cr1^++^/c-Kit^int^ cells (**Figure 3E**, P2) showed an ∼3-fold accumulation of CD115^++^/FLT3^++^ cells in LN and a concomitant 65% depletion in the BM, as early as 3 hours after LPS administration (**Figures 3F-G and Figures S3A-B**). In the absence of significant changes in the LN content of macrophages, these data demonstrate that LPS-induced systemic inflammation results in robust and specific recruitment of phenotypic GMP that are BM cDC committed progenitors to the LN.

### Myeloid progenitor migration to LN is Traf6-dependent and NF-κB-independent

Immune cells and HSC/P express TLR (Beutler and Rietschel, 2003; Nagai et al., 2006; Takeuchi and Akira, 2010) which act as microorganism sensors. LPS stimulation of TLR recruits MyD88 and TRIF through the canonical and endosomal pathways respectively. Both adaptors subsequently recruit TRAF6, which acts as the molecular hub of both signaling branches (Akira et al., 2001; Kawai and Akira, 2006). To determine whether Traf6 deficiency might affect the migration of HSC/P in response to LPS, we exploited an animal model in which *Traf6* is deleted only in hematopoietic cells (Kobayashi et al., 2003) (**Figure 4A, Figure S3C**). LN from *Mx1Cre;WT* chimeric mice after LPS administration (at ZT7, corresponding with the peak of progenitor content in LN, **Figure 1H**) revealed a 3-fold increase in the frequency of CFU-GM. This increase was completely abrogated by the deficiency of Traf6 in hematopoietic cells (*Mx1Cre;Traf6*^Δ/Δ^ animals, **Figure 4B**) indicating that the signals which result in CFU-GM mobilization to LN are mediated by hematopoietic Traf6.

**Figure 4.**
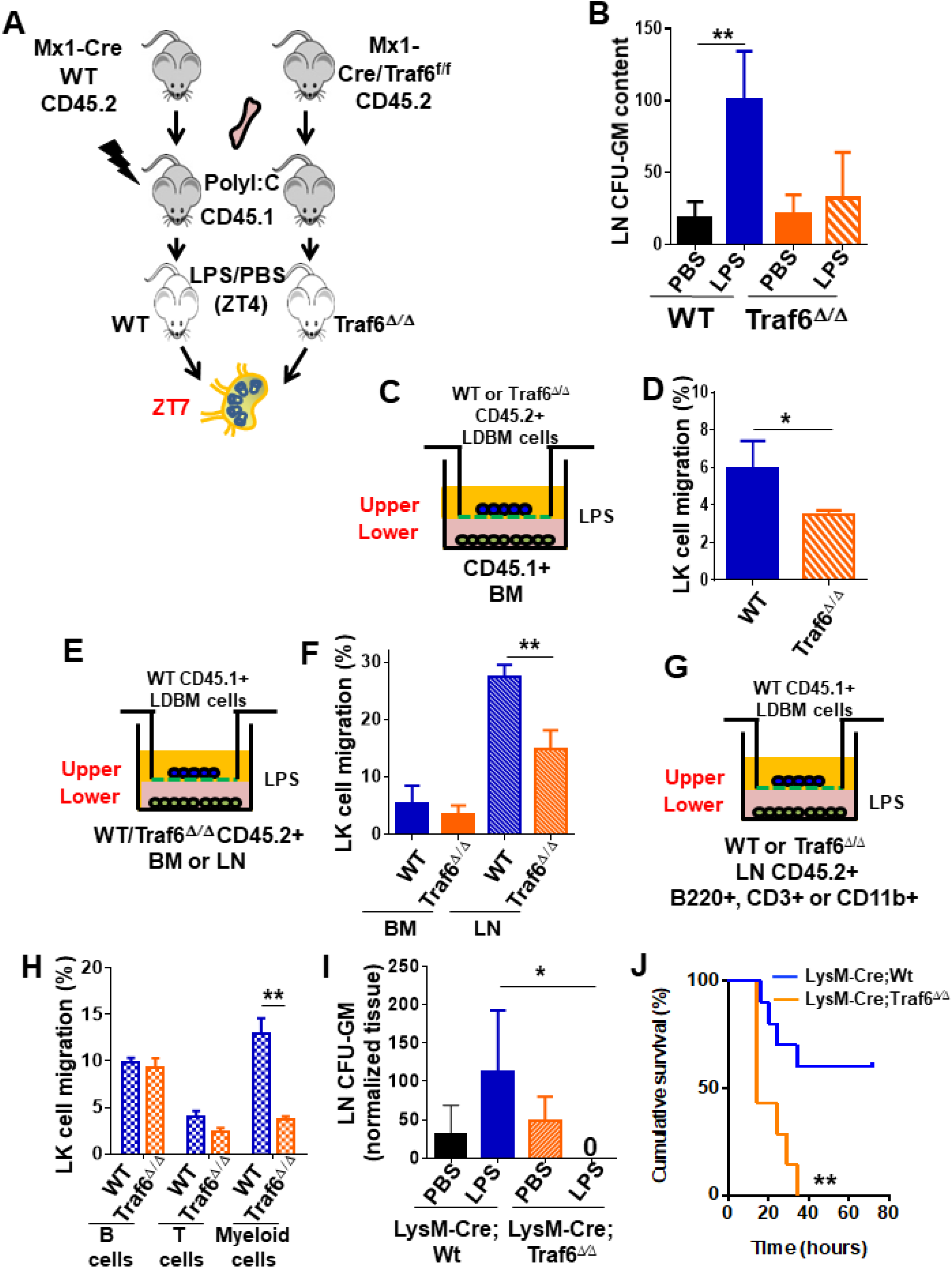
Traf6 is key regulator for migration of BM-derived myeloid progenitors to lymph nodes in a non-cell autonomous manner. (A) Schema of full chimeric mice made by non-competitive transplantation of CD45.2^+^ *Mx1Cre;WT* and *Mx1Cre;Traf6^flox/flox^* BM cells into lethally irradiated CD45.1^+^ B6.SJL^Ptprca Pep3b/BoyJ^. 6 weeks later Traf6 gene were deleted by intraperitoneal injection of poly(I:C). 1 week later we performed PBS/LPS injection early in the rest phase (ZT4 [0h]) and LN-contained myeloid progenitors at ZT7 (3h) was scored by CFU assay. (B) Absolute number of CFU-GM presents in LN from *Mx1Cre;WT (solid bars)* and *Mx1Cre;Traf6^Δ/^*^Δ^ (orange bars) full chimeric mice (n= 6-7 mice per group) after PBS (black and orange solid bars) or LPS (blue and mosaic bars) administration. Values are shown as mean ± SEM of two independent experiments with a minimum of 3 mice per group *P< 0.05, **P<0.01. (C-H) *In vitro* transwell migration assay for BM-derived LK cells. (C) Experimental design for migration of WT or Traf6^Δ/Δ^ low-density (LD)BM cells (CD45.2^+^) toward a WT microenvironment generated by BM (CD45.1^+^) in the presence of LPS for 4 hours. (D) Graph represents the percentage migrated LK from WT (blue solid bar) or Traf6^Δ/Δ^ (orange mosaic bar) low-density BM (LDBM) cells to the bottom as depicted in C. (E). Experimental design for migration of WT LDBM cells (CD45.1^+^) toward a gradient generated by WT/Traf6^Δ/Δ^ (CD45.2^+^) BM or LN cells in the presence of LPS for 4 hours. (F) Graph represents the percentage of LDBM LK migrated to the BM bottom (solid bars) or LN bottom (mosaic bars) as schemed in E. (G) Experimental design for WT LDBM cells (CD45.1^+^) migration toward gradient generated by WT/Traf6^Δ/Δ^ LN-derived T cells (CD45.2^+^/CD3e^+^/CD11b^−^/B220^−^) or B cells (CD45.2^+^/CD3e^−^CD11b^−^/B220^+^) or myeloid cells (CD45.2^+^/CD3e^−^/CD11b^+^/B220^−^) in the presence of LPS for 4 hour. (H) Graph represents the percentage of migrated LK LDBM to the WT LN bottom (blue mosaic bars) or Traf6^Δ/Δ^ LN bottom (orange mosaic bars) as schemed G. In all cases, LK cell migration was determined by CD45 allotype analysis using flow cytometry in triplicate. Values are presented as mean ± SD. (I) Absolute number of CFU-GM presents in LN from *LysM-Cre;WT* (solid bars) and *LysM-Cre;Traf6^Δ/^*^Δ^ (mosaic bars) full chimeric mice after PBS/LPS administration at ZT7 (3h). A minimum of 4 mice per group were analyzed. Values are presented as mean ± SD, *P<0.05. **P<0.01. (J) Graph represent cumulative survival of *LysM-Cre;WT* (blue line) and *LysM-Cre;Traf6^Δ/^*^Δ^ (orange line) after 10 mg/Kg of LPS. **P<0.01. (J) Survival curve after 30 mg/Kg of b.w.injection in *LysM-Cre;WT* (blue line) *or LysM-Cre;Traf6^Δ/^*^Δ^ (orange line). **P<0.01.

Although HSC/P respond directly to PAMPs such as LPS (Nagai et al., 2006; Zhao et al., 2014), direct Gram-negative infection-derived LPS sensing by HSC/P does not play an essential role in emergency granulopoiesis, but rather requires TLR4-dependent signals within the microenvironment (Boettcher et al., 2012; Kwak et al., 2015). To further test whether the hematopoietic Traf6-dependent response to LPS resulting in mobilization of GMP from the BM to the LN by ZT7 is indeed cell autonomous and not determined by the microenvironment, we used the conditional Traf6-deficiency model and analyzed the *in vitro* migration of myeloid progenitors towards chemoattractant gradients generated by LPS-stimulated BM or LN cells in assays designed to identify the hematopoietic cell population affected by LPS (**Figures 4C, E, G**). We found that the cell-autonomous deficiency of Traf6 resulted in a relative decrease in migration of ∼40% (**Figure 4D**) of BM myeloid progenitors in presence of LPS, indicating that Traf6 is required for LPS-dependent cell-autonomous BM myeloid progenitor migration. Interestingly, analysis of non-cell autonomous migration of BM myeloid progenitors demonstrated that LN-derived cells generated more potent chemoattractant signals resulting in a much larger migration of WT BM myeloid progenitors (5-fold higher, ∼30%) during the same period (**Figures 4E-F**), which was drastically diminished (∼50% reduction) by Traf6-deficiency in LN cells, but not when using control BM cells as chemoattractant source (**Figures 4E-F**). These data indicate that although LPS-mobilized myeloid progenitors depend on both cell-autonomous and non-cell autonomous Traf6-dependent signals, the chemoattractant gradient generated by LPS on LN cells is the predominant effect responsible for Traf6-dependent myeloid progenitor migration.

To delineate the resident LN population to cause the migration of GMPs into the LN, we isolated T-cells (CD3e^+^), B-cells (B220^+^) and myeloid cells (CD11b^+^) from LN of WT or Traf6^Δ/Δ^ mice and layered input cell equivalents on the bottom of the chamber with LPS, as in the previous experiments (**Figure 4G**). Although only ∼1% of LN cells are myeloid, we observed that LN CD11b^+^ cells, but not B or T cells, from Traf6^Δ/Δ^ mice can recapitulate the same reduction of progenitor migration achieved by complete LN tissue (**Figure 4H**). To confirm that a Traf6-dependent signaling in LN CD11b+ cells is responsible for myeloid progenitor mobilization and eliminate the possible inflammatory effect of previous treatment with polyI:C in Mx1-Cre Tg mice, we crossed *Traf6^flox/flox^* mice with *LysM-Cre* transgenic mice(Clausen et al., 1999; Cross et al., 1988) and analyzed the migration to LN after LPS administration in mature myeloid lineage-specific Traf6 deficient *(LysM-Cre;Traf6^flox/flox^*) mice. Mature myeloid lineage-specific deletion of Traf6 abrogated the migration of myeloid progenitors to LN in response to LPS (**Figure 4I**). A major consequence of the deficiency of Traf6 in mature myeloid lineage-specific cells was an increase in the endotoxemia dependent mortality (**Figure 4J**) indicating that Traf6 expression in mature myeloid cells is required for both migration of myeloid progenitors to LN and protection of the inflammatory cytokine storm responsible for LPS-induced death. Altogether, these data indicate that LPS/Traf6 signaling is required for migration of myeloid progenitors through predominantly long-range acting, mature myeloid lineage-dependent chemoattractant signals and that LPS/Traf6 signaling in LysM^+^ cells is protective against endotoxin-induced inflammation.

Activation of TLRs conserves inflammatory pathways which culminate in the activation of the NF-κB transcription factors(Karin and Greten, 2005). The LPS binds TLR4/MD2 complexes on the cell surface, and through a series of adaptors and kinases recruits Traf6. By an E3 ligase-dependent mechanism, Traf6 activates the IκB kinase (IKK) complex, which initiates IκBα degradation. Subsequent nuclear translocation of NF-κB transcription factors results in the expression of cytokine and chemokine genes. To determine whether emergent NF-κB signaling is responsible downstream of LPS/Traf6 for the LPS-induced LN migratory effect of myeloid progenitors, we overexpressed a degradation-resistant mutant of IκBα (IκBα super-repressor [IκBα_SR_]), in primary murine progenitors, which were then differentiated into macrophages/cDC by macrophage colony-stimulating factor (M-CSF)(O’Keeffe et al., 2010) (**Figure S3D**). Analysis of LPS-driven migration *in vitro* (**Figure S3E**) demonstrated that the expression of IκBα_SR_ does not reduce the effect of LPS on the migration of myeloid progenitors towards LPS-stimulated macrophages/cDC (**Figures S3F-G**), indicating that NF-κB transcription factors are dispensable for myeloid progenitor migration. In contrast, inhibition of intracellular protein traffic using monensin dramatically decreased myeloid precursor migration (**Figures S3F-G**), suggesting that intracellular protein trafficking is necessary for the migration phenotype. Collectively, these data indicate that LPS-induced myeloid progenitor migration occurs through an NF-κB-independent, intracellular protein traffic-dependent pathway, and suggests that the progenitor mobilizing effect of LPS may not require transcriptional activation, depending rather on the intracellular traffic of secreted proteins.

### Myeloid progenitors home into LN in a CCL19/CCR7 dependent fashion but independently of L-selectin

The secretome of myeloid cells includes multiple cytokines/chemokines with short- and long-range activities on activation, proliferation, survival, differentiation, and migration of target cells. Specifically, secreted chemokines stimulate migration of target cells following chemokines to the areas of highest concentration. It has been described that hematopoietic progenitor migration is dependent on Cxcl12 gradients (Greenbaum et al., 2013; Mendez-Ferrer et al., 2010). However, by ZT7, LPS induced upregulation of Cxcl12 expression in BM, but not in LN, indicating that Cxcl12 tissue concentrations per se could not explain the mechanism of migration to LN (**Figure S4A**). An array of tests on secreted chemokines and cytokines and demonstrated distinct secretome signatures between BM and LN tissues after LPS administration (**Figures S4B-N**). Several myeloid cell cytokines and chemokines with ability to recruit and differentiate macrophages and cDC were found to be upregulated in the extracellular fluid of LN rather than BM as early as one hour after LPS challenge (**Figure S4C-J**). However, none of these candidate cytokines/chemokines were found to consistently generate a differential tissue concentration *in vivo* between LN and BM at both ZT5 and ZT7 (**Figure S4B**). Similar to Cxcl12, some cytokines/chemokines with potential chemoattractant ability were also found to be upregulated in BM rather than in LN or in both tissues similarly (**Figure S4K-N**). The lack of in vivo tissue differential levels strongly suggested that these BM-derived cytokines or chemokines were unlikely to be responsible for the attraction of BM myeloid progenitors to the LN.

The C-C chemokine receptor type 7 (Ccr7) ligand macrophage-inflammatory protein (MIP)-3b/Ccl19 has been reported as a chemoattractant for BM and cord blood CD34^+^ cells *in vitro*, mainly CFU-GM (Kim et al., 1998). Analysis of Ccl19 in the extracellular fluid of the femoral cavity, LN, and plasma, demonstrated that *in vivo* administration of LPS promotes a secretion of Ccl19 in LN when compared with BM and PB (**Figure 5A**). This differential secretion is specific to Ccl19 since Ccl21, a highly-related chemokine, did not show the formation of similar differential tissue concentrations in LN after LPS administration (**Figure S5A**). Ccl19 is secreted by LN myeloid cells after LPS stimulation and depends on Traf6 expression (**Figure S5B**). Ccr7-mediated signals control the migration of immune cells to secondary lymphoid organs such as LN, facilitating efficient surveillance and targeted cellular response (Forster et al., 2008). Also, LPS upregulates membrane Ccr7 expression on cDC and their committed progenitors (Schmid et al., 2011). We therefore hypothesized that LN-trafficking of phenotypic GMP/MDP is regulated by Ccr7, and that therefore the Ccl19/Ccr7 axis might explain the coexistence of cell-autonomous and non-cell autonomous mechanisms required for GMP migration from the BM to LN in response to LPS. To test our hypothesis, we first analyzed whether the specific deficiency of either Ccl19 or Ccr7 modified the level of progenitor migration to LN. To prevent the interference of long-term deficiencies of Ccl19 and Ccr7 expression described in deficient murine models (Forster et al., 1999; Mori et al., 2001), we performed short-term *in vivo* neutralization of Ccl19 ligand or the Ccr7 receptor by using specific antibodies or isotype controls (**Figure 5B**) and determine the content of CFU-GM in LN after LPS challenge or PBS control. We administered an anti-Ccl19 and an anti-Ccr7 neutralizing antibody (or their controls) twice within 15 hours before LPS administration. We found a dramatic reduction (>90%) in the number of CFU-GM in the LN of LPS-, anti-Ccl19 treated animals by ZT7 (**Figure 5C**). Also, we confirmed that Ccr7-expressing GMP in BM rapidly decreased in response to LPS and increased in local LN (**Figures S5D-F**). The abrogation of accumulation of progenitors in LN was reproduced by the administration of anti-Ccr7 (**Figure 5D**). Noteworthy, the administration of anti-Ccr7 phenocopied the effect of Traf6 deficiency in LysM^+^ cells by decreasing the latency to death or increasing the mortality rate of mice treated with lethal (**Figure 5E**) or sublethal (**Figure 5F**) doses of LPS in the first hours after LPS administration, respectively. Second, we analyzed the membrane expression of Ccr7 on BM-derived GMP, CMP and MEP from *Mx1Cre;WT or Mx1Cre;Traf6^Δ/Δ^* mice, with or without LPS stimulation. Membrane Ccr7 levels were significantly upregulated as early as 1 hour after LPS administration on GMP in LPS-treated WT mice. Such upregulation was abrogated in LPS-treated Traf6-deficient GMP (**Figures S5G**). Finally, we confirmed that hematopoietic chimeric Ccl19^−/-^ animals did not mount a migratory response of myeloid progenitors from BM to LN in response to LPS (**Figures 5G-H).**

**Figure 5.**
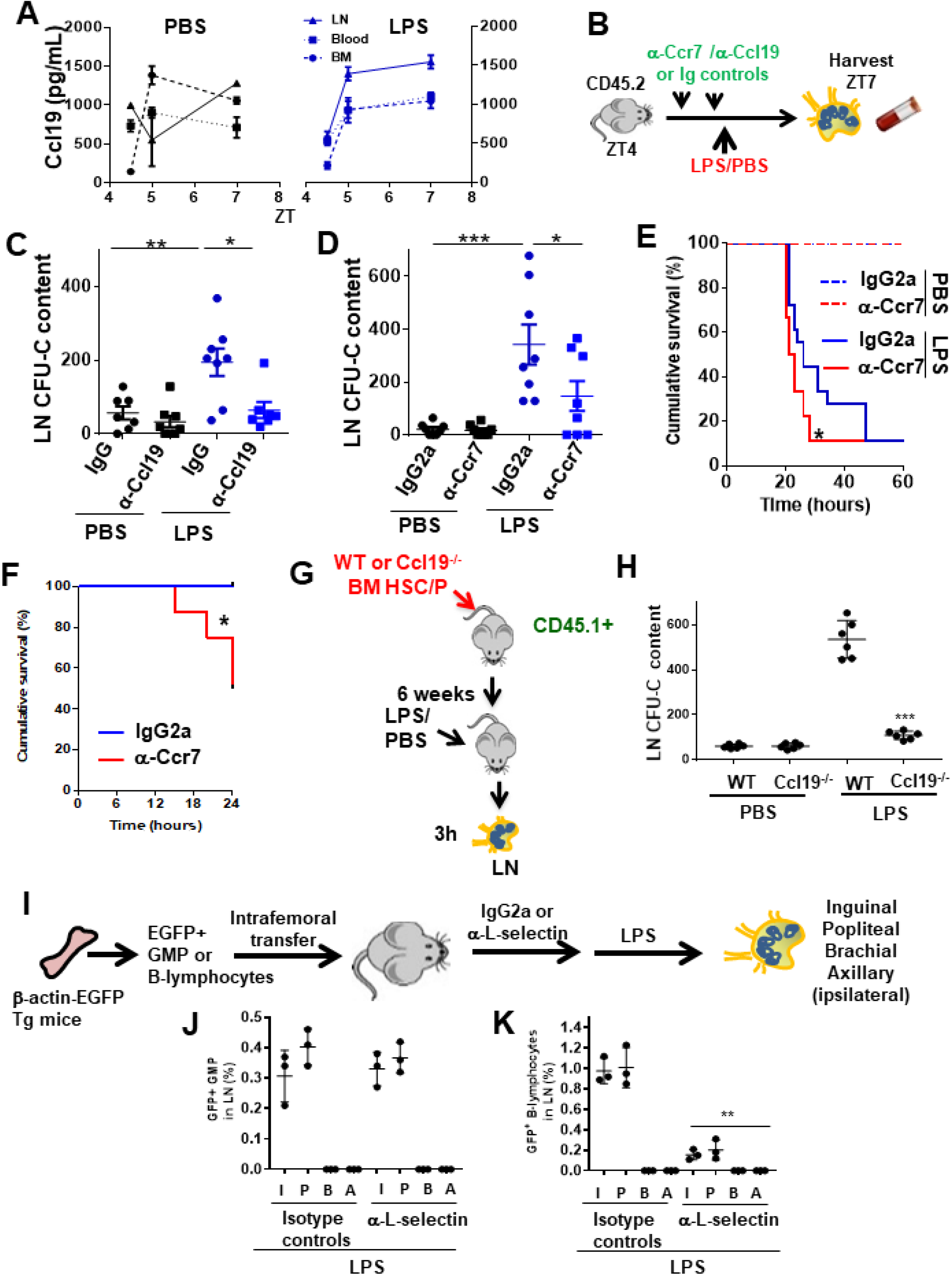
GMP cells drain into local Lymphatics and not blood circulation in early inflammation *via* Ccl19/Ccr7. (A) Graph represents soluble Ccl19 chemokine in femoral or LN extracellular fluid and PB plasma after PBS (black lines) or LPS (blue lines) administration at different circadian cycle times (ZT4.5 [0.5h], ZT5 [1h] and ZT7 [3h]). Values represent mean ± SEM of two independent experiments in duplicate. (B) Strategy for *in vivo* neutralization of Ccr7 receptor or Ccl19 ligand by in2jections of anti-Ccr7 antibody or anti-Ccl19 antibody (50 μg/dose, two doses) into C57Bl/6 mice. One day after the last dose of antibodies, PBS or LPS was administered at ZT4 (0h) and the myeloid progenitors-circulating cells from the LN and PB were measured by CFU assay at ZT7 (3h). (C-D) Absolute number of progenitors present into LN from neutralized mice with anti-Ccl19/IgG (left graph) or anti-Ccr7/IgG2a (right graph) after PBS (black) or LPS (blue) administration as depicted in B (n=7-8 mice per group). Values represent mean ± SEM. *P<0.05 **P<0.01. (E-F) Survival curves after 30 mg/Kg LPS of b.w. (F) and 10 mg/Kg LPS of b.w. or PBS as control (E, dashed lines) into WT C57BL/6 mice pre-treated with anti-Ccr7 (red line) or IgG2a (blue line). *P<0.05. (G) Generation of hematopoietic chimeric Ccl19 expressing (WT) or not (Ccl19^−/-^) mice and isolation of LNs after administration of PBS or LPS. (H) Colony forming unit content of LN from either WT or Ccl19^−/-^ hematopoietic chimeric animals treated with PBS or LPS (***p≤0.001). (I) Experimental design to analyze L-selectin dependence of femoral GMP migration to regional (or distant) LN after LPS administration. (J-K) Percentages of GFP ^+^ cells in LN after administration of an isotype control or anti-L-selectin antibodies. (J) Frequency of GFP ^+^ GMP cells in regional LN after administration of LPS was not modified by L-selectin blockade in vivo. (J) Inhibition of the migration of GFP^+^ B-lymphocytes to regional LNs in mice pre-treated with anti-L-selectin antibody (**p≤0.01). In B and C, LN were collected at ZT7 or 3h after LPS.

Traffic of myeloid progenitors to regional LNs was recapitulated in mice receiving intrafemoral adoptive transfer of GMP (**Figures 5I-J**). In these mice, in vivo L-selectin blockade did not abrogate GMP migration to regional LN while sinusoidal-dependent B-lymphocyte mobilization into regional LN was significantly impaired (**Figures 5J-K)**, indicating that the migration of BM myeloid progenitors, unlike B cells, into the regional lymphatic circulation is L-selectin independent and therefore unlikely to be mediated by LN high endothelial venules (HEV) (Rosen, 2004). Altogether, these data strongly indicated that Ccl19/Ccr7 chemokine signaling is required for the rapid migration of myeloid progenitors to LN upon LPS administration and that Ccr7 signaling is required to prevent death within the first hours after LPS administration. Given the strong time association of these events, these data support a role for the Ccr7-dependent early traffic of myeloid progenitors in the amelioration or delay of the endotoxic shock induced by LPS.

### Ccl19 is expressed and pre-stored in cDC2 and released upon activation of IKK/Snap23

Chemokine secretion requires endosomal fusion with the membrane which can be detected by exposure of the phosphatidylserine (PS)-rich inner leaflet of the endosomes to the external surface of the cell membrane, providing a venue to determine what cell types were responsible for the secretion of Ccl19. We found an increase in the levels of PS residues on the outer membrane leaflet of WT LN cDC2 (defined as CD11b^+^/CD11c^+^), but not in the CD11b^−^ /CD11c^+^ population which comprises cDC1 and plasmacytoid DC (**Figure 6A**). Interestingly, the exposure of PS residues was abrogated in Traf6^Δ∕Δ^ LN cDC (**Figure 6A**) indicating that Traf6 also mediates the process of vesicle secretion upon LPS stimulation. Having demonstrated that LPS/Traf6 signaling is required for chemokine traffic/secretion in LN mature myeloid-lineage cells, and NF-κB transcriptional activation is dispensable for LPS-dependent myeloid progenitor migration, we hypothesized that Traf6 acts through non-NF-kB dependent IKK activity. To determine whether canonical LPS/TLR downstream effectors were involved in the process of myeloid progenitor migration, we analyzed the chemotaxis of BM myeloid progenitors towards a gradient generated by LN cells in the presence of LPS and specific inhibitors for interleukin receptor associated kinase 1/4 (Irak1/4), ubiquitin-conjugating enzyme 13 (Ubc13), and IKKβ (**Figure 6B**). Increased myeloid progenitor migration was reversed by all three specific inhibitors (**Figure 6B**) indicating that the integrity of canonical signaling pathway upstream of NF-κB might be required to attract myeloid progenitors from the BM to the LN. The Traf6/IKK dependent rapid response to LPS strongly suggests that LPS induces secretion of Ccl19 through a mechanism of rapid release from pre-stored pools. The release of pre-formed cytokines in pre-pooled, stored late endosomes depends on IKK activity through the phosphorylation of mediators of cell membrane fusion. SNAP23 is an essential component of the high affinity receptor which is part of the general membrane fusion machinery and an important regulator of transport vesicle docking and fusion (Karim et al., 2013; Suzuki and Verma, 2008). Phospho-SNAP23(Ser95) is significantly upregulated by LPS in LN cDC (**Figure 6C**). Each of the inhibitors for Irak1/4, Ubc13 and IKK abrogated the activation of SNAP23 (**Figures 6C and S5H**). Altogether, this set of data indicates that the activation through Traf6/Irak1/4/Ubc13 induced by LPS activates vesicular fusion and vesicular cargo release of pre-formed Ccl19 accumulated in late endosomes of LN myeloid cells. Analysis of steady-state LN myeloid cell populations identified a subpopulation of cDC but not pDC or macrophages, containing most of the cytoplasmic expression of Ccl19. Further analysis of the subpopulations of LN cDC2 demonstrated that B220^−^/CD8^−^ cDC that expressed the endocytic receptor DEC-205 (CD205^+^) and the mannose receptor signal regulatory protein α (SIRPα, CD172a^+^) distinctly stored high levels of cytosolic Ccl19 (**Figures 6D and S6A)** unlike other DC populations which expressed low levels of Ccl19 (**Figure S6B).** These data indicate that a subpopulation of cDC2 stores intracellular Ccl19 and is potentially able to self-regulate the migration of its own progenitors in inflammation.

**Figure 6.**
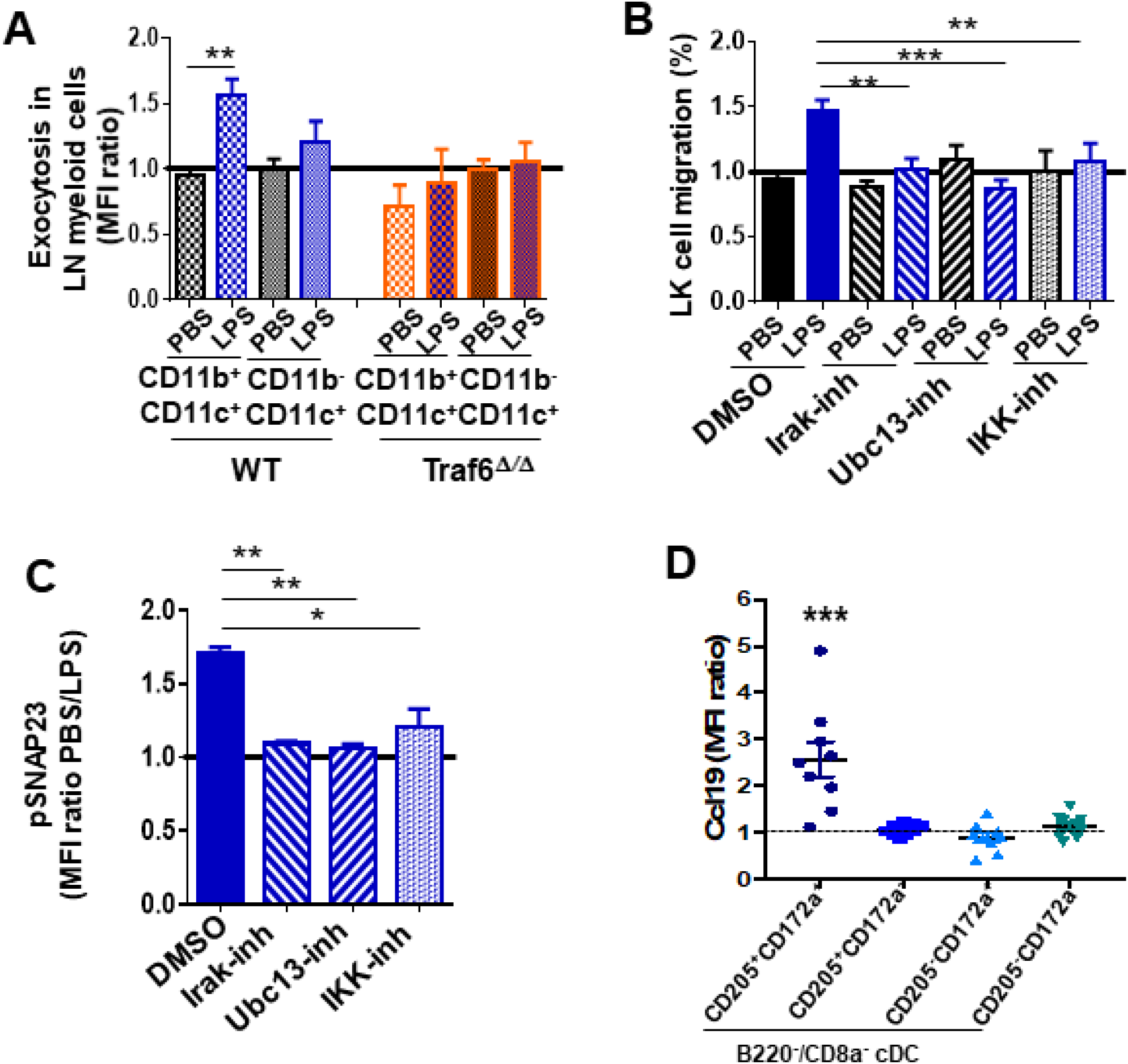
In vivo analysis of Ccl19/Ccr7 axis during inflammation. Pharmacological regulation of LPS/TLR signaling pathway. (A) Annexin-V binding to membrane PS on LN myeloid populations from WT (left mosaic bars) and Traf6^Δ∕Δ^ (right mosaic bars) (n=4 mice per group) after PBS or LPS administration. LN suspension cells were stained for myeloid surface markers including annexin-V and analyzed by flow cytometry. Values represent mean ± SEM. **P<0.01. (B) Transwell migration of LDBM-derived LK cells toward gradient generated by pre-treated LN cells with DMSO (solid bars) as vehicle control and inhibitors (mosaic bars) against Irak1/4 (right lined), Ubc13 (left lined), and IKK (white squares), and following TLR signaling pathway activation by PBS (black) or LPS (blue). (C) Analysis of SNAP23 phosphorylation (Ser95) in LN myeloid cells previously treated with DMSO (solid bar) as vehicle control, Irak1/4 (right lined mosaic bar), Ubc13 (left lined mosaic bar) or IKK (white squares mosaic bar) inhibitors. Values represent two independent experiments as mean ± SEM of two independent experiments performed in triplicate. *P<0.05, **P<0.01. (D) Graph shows intracellular Ccl19 in LN-derived myeloid cells from non-manipulated mice by flow cytometry. Values represent mean ± SEM of three independent experiments, n=8-9. ****P<0.0001.

## DISCUSSION

This study describes a previously unrecognized, rapid, emergent traffic of myeloid progenitor cells from the BM via lymphatic vessels directly to lymphatic tissues that by-pass the peripheral blood stream. Careful analysis of serial femoral sections has not unveiled the existence of a communication between lymphatic and blood vessels in BM further suggesting the lack of communication between both circuits within the BM cavity and thus likely functional regional independence of each circuit. Our data thus also supports the recently described existence of functional lymphatic vessels in the bone. High-resolution confocal and multiphoton microscopy demonstrated the existence of Lyve1+ cells in which their transgenic reporter illuminated upon exposure to high-dose LPS in vivo along with tiny projections of lymphatics penetrating into the bone. Probably, bone processing and cleaning before fixation and decalcification may have deprived us (and other investigators) from a better identification of notable, anatomically identifiable lymphatic vessels within the network of transcortical capillaries (Gruneboom et al., 2019).

Bone is a dynamic organ in constant remodeling. Upon inflammation for example, cytokines and microbial LPS are capable to initiate bone absorption by activating osteoclasts (Hardy and Cooper, 2009; Nason et al., 2009). Systemic inflammation has been associated with osteoclast activation and osteoblast thinning (Hardy and Cooper, 2009; Nason et al., 2009) and bone lymphatic endothelial cells have been shown to arise rapidly from pre-existing regional lymphatics upon osteoclast activation (Hominick et al., 2018; Monroy et al., 2020). Osteoclast activation and osteoblast thinning are likely to facilitate transcortical migration of cells and fluid through existing transcortical vessels.

Our data showed that as early as 90 minutes after LPS administration, myeloid progenitors to or are in closer proximity to the lymphatic endothelium in BM while 1.5 hours later there is a ∼70% reduction of myeloid progenitors within the BM and a marked increase of myeloid progenitors in the LN tissue. This observation along with the need of a longer period of time to detect an increase in the frequency of myeloid progenitors in peripheral blood suggests that two temporally distinct waves of progenitors take place, a fast one to the lymphatic circulation followed by a slower one into the blood stream.

In our study, by using transgenic animals, we demonstrated that the administration of a single dose of LPS suffices to induce migration of GMP/MDP while no other types of progenitors or stem cells migrate to lymphatics in this first wave of egression before any significant contribution from or to PB and replenish short-lived cDCs in murine acute model of inflammatory signaling by LPS, and that therefore they may modulate the course of infectious diseases and other inflammatory conditions. This traffic is also likely to happen in homeostatic conditions as previously shown (Waskow et al., 2008) while our analysis provides compelling evidence on its striking activation upon inflammation/LPS administration.

Our data support the migration of a distinctly immature progenitor population composed of GMP/MDP with ability to generate cDC in LN upon traffic from BM to LN. This traffic of myeloid progenitors from BM to LN can be recapitulated in mice receiving intrafemoral adoptive transfer of GMPs. These GMP/MDP tend to localize in the T-cell areas of LN. Cheong et al. reported that migratory monocyte-derived cDC2 can also localize in T cell areas of the LN and acquire an inflammatory phenotype DC-SIGN/CD209a^+^ (Cheong et al., 2010). No significant mobilization of M-CSF responding monocyte progenitors (CFU-M) can be found as early as 3 hours after LPS administration. Interestingly, the interference of this traffic by either blocking the chemokine secretion by mature LysM-expressing myeloid cells, or by blocking the chemokine receptor Ccr7 results in increased animal death, which strongly suggests that the traffic of progenitors resulting from Ccl19/Ccr7 signaling is not only destined to the differentiation into cDC but also to a more immediate anti-inflammatory role as suppressors of the endotoxic shock effects. Ccr7^+^ GMP/MDP, but not other myeloid or lymphoid progenitors, egress BM. Such egress follows differential tissue levels of Ccl19 resulting from activation of the secretion of the pre-formed chemokine by LN LysM-expressing mature myeloid cells, specifically a subpopulation of cDC expressing CD205 and CD172a. This process seems to be independent of Cxcl12 levels since no changes in Cxcl12 levels in LN, PB or BM were observed and this effect seems to be exclusively dependent on activation of non-canonical Traft6/IKK activity without need for transcriptional activation. Schmid et al (Schmid et al., 2011) demonstrated that a population of common dendritic progenitors (CDP), a non-GMP derived population of progenitors can also migrate from the BM to lymphoid and non-lymphoid tissues in response to TLR agonists and generate both cDCs and pDCs (pDCs). The type of migration though depended on combined downregulation of Cxcr4 and upregulation of Ccr7, which seem to imitate the mechanism of GMP migration. Interestingly, Ccl21, which is expressed by lymphatic endothelial and stromal cells but not by myeloid cells (Eberlein et al., 2010), does not induce any differential gradient of secretion between LN and BM or blood suggesting that Ccl21 may not be a primary mediator of the myeloid progenitor migration from BM to LN upon LPS challenge, while the hematopoietic deficiency of Ccl19 suffices to completely abrogate the mobilization of myeloid progenitors to LN induced by LPS.

Our data support the existence of a steady-state LN population of cDC which co-expresses the maturation antigens CD205 and CD172a and store high levels of Ccl19 in their cytoplasm. An interesting possibility is, as our data indicate, that upon bacterial antigen challenge, differentiated myeloid cells of LN like cDCs, which respond to LPS by secreting chemokine-containing pre-formed exosomes, accelerate a positive feedback activation loop to recruit cDC progenitors to the lymphatic tissue. cDC in LNs might thus act as sensors for the presence of bacterial products and release Ccl19 within minutes. Individual DCs have a short half-life (1.5-2.9 days)(Kamath et al., 2002) and DC precursors have a short half-life in blood circulation (Breton et al., 2015). We posit that the migration of DC progenitors through the lymph tissues provides a direct afferent communication between the LN mature cDC population responsible for the secretion of the chemokine Ccl19 and at the same time, allows the emergent migration of functional cDC progenitors from the BM to replenish the repertoire of lymphatic antigen presenting cells.

Finally, our data also support the key role of an alternative inflammatory signaling pathway elicited by coordinated by Traf6/IKK responsible for SNAP23 phosphorylation and Ccl19 secretion, before resulting in transcriptional regulation by their downstream effector NF-kB. Traf6 has been identified as a signaling molecule that can regulate splicing of downstream targets without affecting NF-kB in hematopoietic stem cells and progenitors (Fang et al., 2017). Our data further identifies non-canonical signaling pathways elicited by Traf6 in differentiated myeloid cells to modulate the inflammatory response affecting the circulatory dynamics of hematopoietic progenitors. Traf6 dependent, cytosolic mediated inflammatory response allows a fast response before inflammatory transcriptional and post-transcriptional signatures are mounted.

In summary, we describe, upon inflammation, a rapid trafficking of cDC biased myeloid progenitors from the BM, via lymphatic vessels, directly to lymphatic tissues that by-passes the blood stream. This GMP/MDP migration represents a mechanism for fast replenishment of cDCs in lymphatic tissues. Rapid replenishment of cDC-biased progenitors in LN may represent a major homeostatic function of this novel lymphatic circuit and may explain why the circulation of myeloid progenitors is conserved during the postnatal life.

## MATERIAL AND METHODS

### Mice

CD57Bl/6 (CD45.2^+^) mice were used between 8-10 weeks of age and were purchased from Jackson Laboratory, Bar Harbor, ME; Harlan Laboratories, Frederick, MD. *Mx1Cre^+^;Traf6*-floxed mice were generated by breeding Mx1-Cre transgenic mice(Mikkola et al., 2003) with biallelic TRAF6 floxed mice (kindly provided by Dr. Yongwon Choi, University of Pennsylvania)(Kobayashi et al., 2003). Full chimeric mice were generated by non-competitive transplantation of *Mx1Cre^Tg/+^;*WT or *Mx1Cre;Traf6^flox/flox^* whole BM cells into lethally irradiated B6.SJL^Ptprca Pepcb/BoyJ^ (CD45.1^+^) mice obtained from the CCHMC Animal Core. Traf6 was deleted upon induced expression of Cre recombinase after 3-6 intraperitoneal injections (10 mg/Kg/b.w. Poly(I:C); Amersham Pharmacia Biotech, Piscataway, NJ, USA) every other day at 6 weeks after transplantation. LysM-Cre;Traf6 floxed mice were generated by non-competitive transplantation of *LysMCre^Tg/+^*;WT or *LysMCre^Tg/+^; Traf6^flox/flox^* whole BM cells into lethally irradiated B6.SJL^Ptprca Pepcb/BoyJ^ (CD45.1^+^) mice obtained from the Division of Experimental Hematology/Cancer Biology of Cincinnati Children’s Hospital Research Foundation (CCHRF).

Lyve1-eGFP (Pham et al., 2010) and β-actin-eGFP (Okabe et al., 1997) and Cx3cr1-GFP(Jung et al., 2000) transgenic mice were purchased from Jackson Laboratories. C57BL/6 mice for circadian cycle analysis of CFU-C were maintained on a 14-hour light / 10-hour darkness lighting schedule. This study was performed in strict accordance with the recommendations in the Guide for the Care and Use of Laboratory Animals of the National Institutes of Health. All of the animals were handled according to approved institutional animal care and use committee (IACUC) protocol #2019-0041 of Cincinnati Children’s Hospital.

### Human specimens

Lymphadenopathies from patients were obtained through Institutional Review Board-approved protocols of the Hospital Reina Sofia (Cordoba, Spain), donor informed consent and legal tutor approval in the case of patients younger than 18 years old. Diagnostic lymphadenopathy biopsies from sixty-four consecutive patients from 2009 until 2013 were analyzed in this study. The median age of patients was 34 years old (range: 3-89). Diagnosis and histological classification of the type of lymphadenopathy and tumors were based on previously published criteria (Campo et al., 2011; Weiss and O’Malley, 2013). Anatomical location of lymphadenopathies is described in Table 1. Specimens were blindly analyzed through adjudication of unique identifiers.

### LPS injection and samples collection

Mice received a single intraperitoneal injection of 30 mg per Kilo of E. coli LPS (Sigma-Aldrich, St Louis, MO) or PBS as vehicle control and were executed always at ZT4 or 4 hours after the initiation of light into the animal room. At different time points after PBS or LPS administration, BM cells from femurs, tibias and pelvis were harvested by crushing in PBS containing 2% of FBS and erythrocytes were lysed using a hypotonic buffer from BD Biosciences. Blood was collected by retro-orbital bleeding or cardiac injection. Liver, kidney and thoracic duct cells were harvested by enzymatic digestion solution with collagenase II (1 mg/mL, ThermoFisher Gibco, Waltham, MA) and dispase (5 mg/mL, Gibco, Life Technologies) in a shaking water bath at 37^0^C for 1h. Spleen cells were isolated by scraping with slides in sterile PBS following red blood cell (RBC) lysis (Pharm Lyse^TM^; BD Bioscience, San Jose, CA). Extracted LN were derived from the cervical and axillary chains exclusively.

### Myeloid progenitor counts assay *in vitro*

Cells from BM, lymph nodes, thoracic duct, blood, spleen, liver and kidneys were depleted of RBC by 2 minute-incubation in Pharm Lyse^TM^ (BD Biosciences, San Jose, CA), washed, counted and plated in semisolid methylcellulose media (Methocult 3434; StemCell Technology, Vancouver, Canada) and cultured in an incubator (37°C, 5% CO_2_/>95% humidity) and the number of CFU-C was scored on day 7 or 8 of culture using an inverted microscope. To examine the type of myeloid progenitors migrating into LN, we used base methylcellulose medium (Methocult 3134; StemCell Technology, Vancouver, Canada) supplemented with 30% FCS, 1% protease-free, deionized BSA (Roche), 100 mM b-mercaptoethanol, 100 IU/mL penicillin, 0.1 mg/mL streptomycin andany of the following: for CFU-GM, rm-GM-CSF (100ng/mL, PeproTech Rocky hill NJ) for specific analysis of CFU-GM content, rm-M-CSF (100ng/mL, PeproTech Rocky hill NJ) for specific analysis of CFU-M content or rh-G-CSF (100ng/mL, Neupogen) for specific analysis of CFU-G content.

### Long-term competitive repopulation assay

To analyze the long-term reconstitution capacity of HSC/Ps mobilized into LNs after LPS administration at different time of periods, 4-5×10^6^ of erythrocyte-depleted CD45.2^+^ LN suspension cells were prepared in sterile conditions and transplanted together with 2.5×10^5^ CD45.1^+^ BM competitor cells into lethally irradiated CD45.1^+^ recipient mice. In some experiments, 10^4^ LN SP cells or 10^3^ BM SP cells from CD45.2^+^ mice were competitively transplanted into CD45.1^+^ recipient mice. Competitive repopulating units (CRU) analysis was performed by flow cytometry analysis (BD Biosciences) at different time points post-transplantation(Harrison, 1980).

### Flow cytometry analysis and cell sorting

For immunophenotype analysis of HSC/P populations by fluorescence-activated cell sorter (FACS), erythrocyte-depleted BM cells were stained first for lineage markers with biotin-labeled mouse lineage panel (BD Biosciences, Pharmingen) containing anti-CD3e (CD3_Ɛ_ chain), anti-TER-119/Erythroid cells (Ly-76), anti-Gr1 (Ly6G and Ly-6C), anti-CD45R (B220), anti-CD11b (integrin α chain, Mac1α) followed of allophycocyanin and cyanine dye Cy7-(APC-Cy7) conjugated streptavidin, allophycocyanin (APC)-conjugated anti-c-Kit (clon 2B8), R-phycoerythrin and cyanine dye Cy7 (PECy7)-conjugated anti-Sca1 (clone D7), eFluor 450-conjugated anti-CD34 (clone RAM34) (affymetrix eBioscience, San Diego CA), PerCP and cyanine dye Cy5.5 (PerCP Cy5.5)-conjugated anti-Fcγ-RII/III (clone 2.4G2) (BD Biosciences). FACS sequential discrimination on a Lineage negative gated population was used to identified LK myeloid progenitor subpopulations: Lin^−^,c-Kit^+^Sca1^−^CD34^+^ FcγRII/III^+^ (granulocyte-macrophage progenitors, GMP); Lin^−^,c-Kit^+^Sca1^−^CD34^+^FcγRII/III^lo^ (common myeloid progenitors, CMP); Lin^−^ cKit^+^Sca1^−^ CD34^+^ FcγRII/III^−^. LSK (Lin-Sca1+c-Kit-) subpopulations were distinguished as Lin^−^,c-Kit^−^Sca1^+^ CD34^−^Flt3^−^ for LT-HSC Lin^−^,c-Kit^−^Sca1^+^CD34^+^Flt3^−^ for ST-HSC and Lin^−^,c-Kit^−^Sca1^+^CD34^+^Flt3^+^ for multipotent progenitors (MPP). For chimera analysis in repopulated animals, 20 µL of red cell-depleted blood was stained with fluorescein isothiocyanate (FITC)-conjugated anti-CD45.1 (clone A20), R-phycoerythrin and cyanine dye Cy7 (PECy7)-conjugated anti-CD45.2 (clone 104), allophycocyanin (APC)-conjugated anti-CD11b (clone M1/70), allophycocyanin and cyanine dye Cy7-(APC-Cy7)-conjugated anti-B220 (clone RA3-6B2), R-phycoerythrin (PE)-conjugated anti-CD3e (clone 145-2C11) and BD Horizon™ V450-conjugated anti-Gr1 (clone RB6-8C5), PerCPefluor^®^710 anti-CD115 (clone AFS98) and R-phycoerythrin conjugated anti-CD135 or anti-Flt3 (clone A2F0.1). All monoclonal antibodies were purchased from BD, Pharmingen. Cell acquisition was performed by flow cytometry (LSRFortessa I, BD Biosciences) equipped with FACSDIVA^™^ software (BD, Biosciences) for multiparameter analysis of the data. FACS sorting strategies were: CD45.1^−^ CD45.2^+^CD3_Ɛ_^+^B220^−^CD11b^−^ for LN-T cells, CD45.1^−^ CD45.2^+^CD3_Ɛ_^−^B220^+^/CD11b^−^ for LN-B cells and CD45.1^−^ CD45.2^+^CD3_Ɛ_^−^/B220^−^/CD11b^+^ for LN-myeloid cells in a FACSAria II cell sorter (BD Biosciences). For BM and LN -side population (SP) cells analysis and sorting, 2×10^6^ cells/mL were stained with Hoescht 3342 (5 µg/mL) as described previously^15^. For intracellular analysis of the phosphorylated state of SNAP23 protein, surface antigen-labeled cells were fixed with Cytofix buffer (BD Biosciences) for 20 min and then permeabilized using Cytofix/Cytoperm buffer (BD Bioscience) for 20 minutes. After washing, cells were stained intracellularly using a rabbit non-conjugated monoclonal anti-phospho-SNAP23(Ser^95^)(Karim et al., 2013) for 40 minutes in Perm/Wash Buffer 1x (BD Bioscience) with 0.5% of rabbit serum. Cells were then incubated with a secondary Alexa Fluor® 488-conjugated (ThermoFisher-Invitrogen) goat anti-rabbit antibody for 40 minutes in Perm/Wash Buffer 1x with 0.5% of goat serum. All incubations after cell stimulation were done on ice and in darkness. Single cell analysis was performed by flow cytometry and the histogram-overlay graphed (LSRFortessa I; FlowJo xV0.7 software; BD Biosciences). The mean fluorescence intensity (MFI) ratio was calculated as the ratio of the fluorescence intensities of LPS-stimulated to PBS-stimulated (control).

### Annexin-V binding

LN suspension cells from *Mx1Cre;WT and Mx1Cre;Traf6^Δ/Δ^* mice were obtained to performed LPS stimulation. 10^6^ cells were plated into 24-well-plates and treated with PBS or LPS for 1 hour. After 15 minutes labelling with surface antibodies against CD45.2 (Clone 104), CD11c (clone HL3), CD11b (Clone M1/70) and B220 (clone RA3-6B2) the samples were washed twice and then stained for annexin-V for 15 minutes and in darkness. All antibodies we purchased from BD Biosciences - Pharmingen. Single cell analysis was performed using flow cytometry and the histogram-overlay graphed (LSRFortessa I; FlowJo xV0.7 software; BD Biosciences). The MFI ratio between LPS MFI and PBS MFI was calculated.

### Homing and seeding assays

For homing assays, 2×10^6^ of Lin^−^ cells, previously depleted by immunomagnetic selection (Lineage Cell Depletion kit, Miltenyi Biotec, Auburn CA), were stained by 1,1^’^-dioctadecyl-3,3,3^’^,3’, tetramethylindocarbocyanine perchlorate;CILC18(3) (5 µM/mL DiI, ThermoFisher-Invitrogen) and adoptively transferred intravenously into non-myeloablated Lyve1-GFP+ mice. Seventeen hours later, one single LPS dose (3mg/mL) or vehicle control (PBS) was administered intraperitoneally. 3 and 6 hours later (ZT7 or ZT10) mice were euthanized with pentobarbital (60-80 mg/Kg) and the whole body was fixed using a freshly made solution of PBS plus 2% of paraformaldehyde (PFA) and 0.05% of glutaraldehyde infused by perfusion pump through left ventricle of the animal. 15-20 minutes later the BM cells and LN organs were harvested and the percentage of labeled Lin^−^ cells which had homed into BM was determined by FACS analysis. The homing calculation was done as previously reported(Boggs, 1984).

### Microscopy

Fixed LN organs were permeabilized for 15 minutes with PBS containing 0.2% of Triton X-100. To detect GFP on the lymphatic endothelium, LN were incubated overnight with a primary antibody anti-GFP+ (ThermoFisher-Invitrogen). LN were scanned by confocal microscopy (Nikon A1R GaAsP) through multidimensional acquisition to construct 3D representations of the whole organ at 10x magnification. The merged images of GFP/DiI or DAPI/DiI are presented and the total cell number of labeled Lin^−^ cells was counted manually. Finally, harvested femurs were decalcified for 14 days with 10% of EDTA (Sigma-Aldrich St Louis, MO) in PBS and embedded in paraffin. Longitudinal section of bone were cut 4-μm thickness and then were des-paraffinized and broke the protein cross-link before stain by antigen retrieval treatment with citrate buffer pH 6 (Cancelas et al., 2005). Then bone sections were permeabilized with 0.2% of Triton X-100 for 15 min and blocked with 5% of BSA for 1h. Slides were stained with primary antibodies anti-GFP (chicken polyclonal, Abcam Inc. Cambridge MA) and rat anti-mouse panendothelial cell antigen (clone, MECA-32, BD Pharmingen) at 4°C overnight. Then we stained with secondary antibodies; goat anti-rat Alexa Fluor-488 and goat anti-chicken Alexa Fluor-568, all from Invitrogen at 1:1000v/v concentration for 1h at room temperature. Blood and lymphatic vessels were scanned by confocal microscopy (Nikon A1R GaAsP) through multidimensional acquisition to construct 3D representation.

To further characterized lymphatic system in bone tissue and BM cavity and to image the close proximity of homed Lin^−^/DiI^+^ to lymphatic vessels into Lyve1-eGFP^+^ mice, we utilized multi-photon intravital microscopy (IVM) as previously described (Gonzalez-Nieto et al., 2012; Kohler et al., 2009). After LPS/PBS injections long bones were harvested and muscle were carefully cleaned. Further, bone tissues were cautiously trimmed with an electric drill (Dremel) to get better excess of the BM cavity for imaging by leaving a very thin (∼30-40 μm) layer of bone tissue. Bones were mounted in 2% low melting agarose to minimize movements during imaging and covered with PBS. Multi-photon microscopy on the long bones (femur and tibia) was subsequently performed using a Nikon A1R Multiphoton Upright Confocal Microscope equipped with Coherent Chameleon II TiSapphire IR laser, tunable from 700 to 1000 nm and signal was detected by low-noise Hamamatsu photomultiplier (PMT) tubes. Bone tissue was identified as second-harmonic (SHG) signal (PMT). Bones were images in PBS using a 25X Apo 1.1 NA LWD water Immersion objective and NIS image software. For Initial standardization, bones were scanned at wavelength of 800, 850, 900 -nm detecting GFP (530 nm) and Dii red (580 nm). For imaging, a 500 × 500-μm area was scanned in ∼35 steps of 4 μm down to 120-150 μm depth using an illumination wavelength of 800 nm detecting SHG signal (480 nm), green (530 nm) and red (580 nm) fluorescence. Control C57BL/6 mice were used as a negative control for Lyve-1 GFP mice to detect specific signal for GFP-lymphatic system in bone tissue and bone marrow cavity. Lymphatic vessels were well detected in the bone tissue using Lyve-1 GFP mice with DiI labeled Lin^−^ cells in the BM cavity. For quantification of proximity of Lin^−^/DiI^+^ with Lymphatic system, Imaris software was used to measure distance between DiI labeled cells and GFP positive lymphatic vessels using 3D images.

### Analysis of L-selectin dependence of femoral GMP/MDP migration to regional LN

C57Bl6 mice received single intraperitoneal injections of MEL14 (CD62L) antibody (Biox-cell) 200ug. Control mice received same amount of Rat IgG2a. Post 3 days of MEL14 antibody treatment, LDBM cells from ß-actin eGFP mice were injected intrafemorally. LPS (30mg/kg, BW) was injected post 1hr. of interfemoral injections. Mice were sacrificed at 3 hours post-administration and ipsilateral and contralateral regional and distant LNs (inguinal, popliteal, axillary and cervical) were isolated for analysis. LN cells were stained GMP markers and anti-CD19-PECy7 (Cat#552854, BD Biosciences) and analyzed by flow cytometry and quantified the GFP^+^ GMP and B lymphocyte populations migrating to LN.

### Femoral GFP^+^ progenitor migration in WT and Ccl19^−/-^ hematopoietic chimeras

Hematopoietic chimeras of WT and Ccl19^−/-^ (Link et al., 2007) BM cells were generated by transplantation into CD45.1+ mice, similarly to Mx1Cre;Traf6^f/f^ hematopoietic chimeras. Mice were followed for 8 weeks and found to have >95% chimera of CD45.2+ cells in peripheral blood. After 8 weeks, femoral LDBM cells from donor congenic -actin transgenic, CD45.2+ mice were injected (5×10^5^ per mouse) intrafemorally to both Wt and Ccl19 hematopoietic chimeric mice. PBS or LPS (30mg/Kg. b.w.) were injected at 1 h post-intrafemoral injections and sacrificed at 3 h post-administration of LPS. At that time, ipsilateral LN from inguinal and popliteal regions were isolated. Suspension of LN cells were counted and stained with specific antibodies for GMP and MDP characterization, and the frequency of different GFP^+^ GMP and MDP populations was analyzed by flow cytometry as mentioned above.

### Chemotaxis/Migration assays

For non-cell autonomous effect analysis, 5×10^5^ of BM or LN nucleated cells from *Mx1Cre^+^;WT or Mx1Cre^+^;TRAF6^Δ/Δ^* CD45.2^+^ were layered on bottom wells of 24-well transwell plate (Corning Inc., Lowell, MA) together with 100 ng/mL of LPS, and 1×10^5^ WT CD45.1^+^ LDBM cells were layered on upper chamber at 37°C, 5% CO_2_. For cell autonomous effect analysis, 5×10^5^ of BM or LN nucleated cells from WT CD45.1^+^ mice were layered in the lower chamber with 100 ng/mL of LPS, and 1×10^5^ *Mx1Cre^Tg/+^*;WT or *Mx1Cre^Tg/+^; Traf6*^Δ/Δ^ CD45.2^+^ LDBM cells were layered in the upper chamber at 37°C, 5% CO_2_. After 4 hours, cells were resuspended and those adhered to the bottom layer were collected using an enzyme free cell dissociation buffer (Cell Dissociation Buffer, enzyme free, PBS, ThermoFisher-Gibco). Progenitor responses toward migratory gradient were analyzed by flow cytometry analysis of LK cells. The percentage of migration were calculated by dividing the number of LK in the outputs by the number of LK in the inputs and multiplied by 100. PBS was included as negative control. All assays were performed in triplicate.

### NFκB activity repression in myeloid cells

To analyze the NFκB-dependent or independent mechanism of myeloid progenitor migration, BM Lin^−^ were transduced with pMSCVpuro-eGFP bicistronic retroviral vector encoding the full length of IκBα mutant (super-repressor) in the presence of the recombinant fragment of fibronectin, CH296 (Takara Bio Inc, Madison, WI) for 16 hours at 37°C. 24 hours later GFP^+^ cells were sorted and macrophages were generated(Chang et al., 2014). To characterize the expanded population, R-phycoerythrin (PE)-conjugated anti-CD169 (clone 3D6.112), PerCP-efluor^®^ 710-conjugated anti CD115 (clone AF598) (affymetrix eBioscience), allophycocyanin and cyanine dye Cy7-(APC-Cy7) conjugated anti-CD11b, efluor^®^ 450-conjugated anti-F4/80 (clone BM8) (affymetrix eBioscience), and Alexa Fluor^®^ 647-conjugated anti-CD68 (clone FA-11)(BD Biosciences) were used for FACS analysis. 50×10^3^ differentiated and transduced macrophages with empty or IκBα super-repressor were layered on bottom wells of 24-well transwell plate in presence of 100 ng/mL of LPS and 1×10^5^ WT LDBM cells were layered on upper chamber at 37°C, 5% CO_2_. Four hours later migrated LK cells were determined by flow cytometry as described above. All assays were performed in triplicate.

### Secretome and individual cytokine/chemokine analyses

BM, plasma and LN cells were isolated in PBS containing a protease inhibitor cocktail (Roche Diagnostics, Chicago IL) and Ccl19/Ccl21 levels were determined by indirect sandwich of enzyme-linked immunosorbent assay (ELISA) following manufacturer’s instructions (**R&D Systems, Minneapolis, MN).** Multi-analytic profiling beads using Milliplex® Multiplex mouse cytokine/chemokine panel I kit (EMD Millipore, Billerica, MA) according to manufacturer’s instructions were used to analyze chemokines and cytokines profile into BM and LN tissues at different time periods after LPS or PBS administration into WT mice.

### In vivo administration of anti-Ccr7 and anti-Ccl19

Monoclonal rat IgG2a antibody specific for Ccr7 (Clone 4B12) or polyclonal goat IgG antibody for Ccl19 (AF880) and and control rat IgG2a or control goat purified IgG, were obtained from R&D Systems. Fifty μg of antibodies were injected twice into C56BL/6 mice within 15 hours (first dose i.v. and the second dose i.p.)

### Small molecule inhibitors

The Irak1/4 Inhibitor I, ubiquitin-conjugating enzyme E2 N (UBE2N) inhibitor, Ubc13 inhibitor (Rhyasen et al., 2013) and IκB Kinase inhibitor (PS-1145 dihydrochloride) were purchased from Sigma-Aldrich. LN cells from C57BL6 mice were treated with 10μM of IRAK-Inh, 0.2 μM of Ubc13-Inh and 10μM of IKK-Inh for 45 minutes and compared with the vehicle dimethylsulphoxide (DMSO) at 0.1% in PBS. Monensin (eBioscience) was used at 2 µM.

### Statistical analysis

Quantitative data is given as mean ± standard error of the mean (SEM) or standard deviation (SD). Statistical comparisons were determined using an unpaired Student t-test, non-parametric Mann-Whitney test, one-way or two-way Anova with Bonferroni corrections. A value of p< 0.05 was considered to be statistically significant.

## ACKNOWLEDGEMENTS

We want to thank Dr. Andre Olsson for flow cytometry analysis advice and Ms. Margaret O’Leary for editing manuscript. We also want to thank Dr. Andres Hidalgo (CNIC, Madrid, Spain and Yale University, New Haven, CT), Drs. Daniel Lucas, Leighton Grimes, Michael Jordan, Edith Jansen and Jizhou Zhang (Cincinnati Children’s Hospital Medical Center) for helpful discussions on data interpretation and experimental approaches. This project has been funded by the Junta de Andalucía of Spain (J.S-L) and NIH R01 GM110628 and DK124115 grants (JAC). The authors declare they have no relevant conflicts of interest. We want to thank Francisco M. Gutierrez for human CFU-C pictures. We also want to thank Jeff Bailey and Victoria Summey for technical assistance and the Mouse and Research Flow Cytometry Core Facilities, both supported by the NIH/CEMH grant P30DK090971-01.

## Supplementary Figure Legends

**Figure S1.**
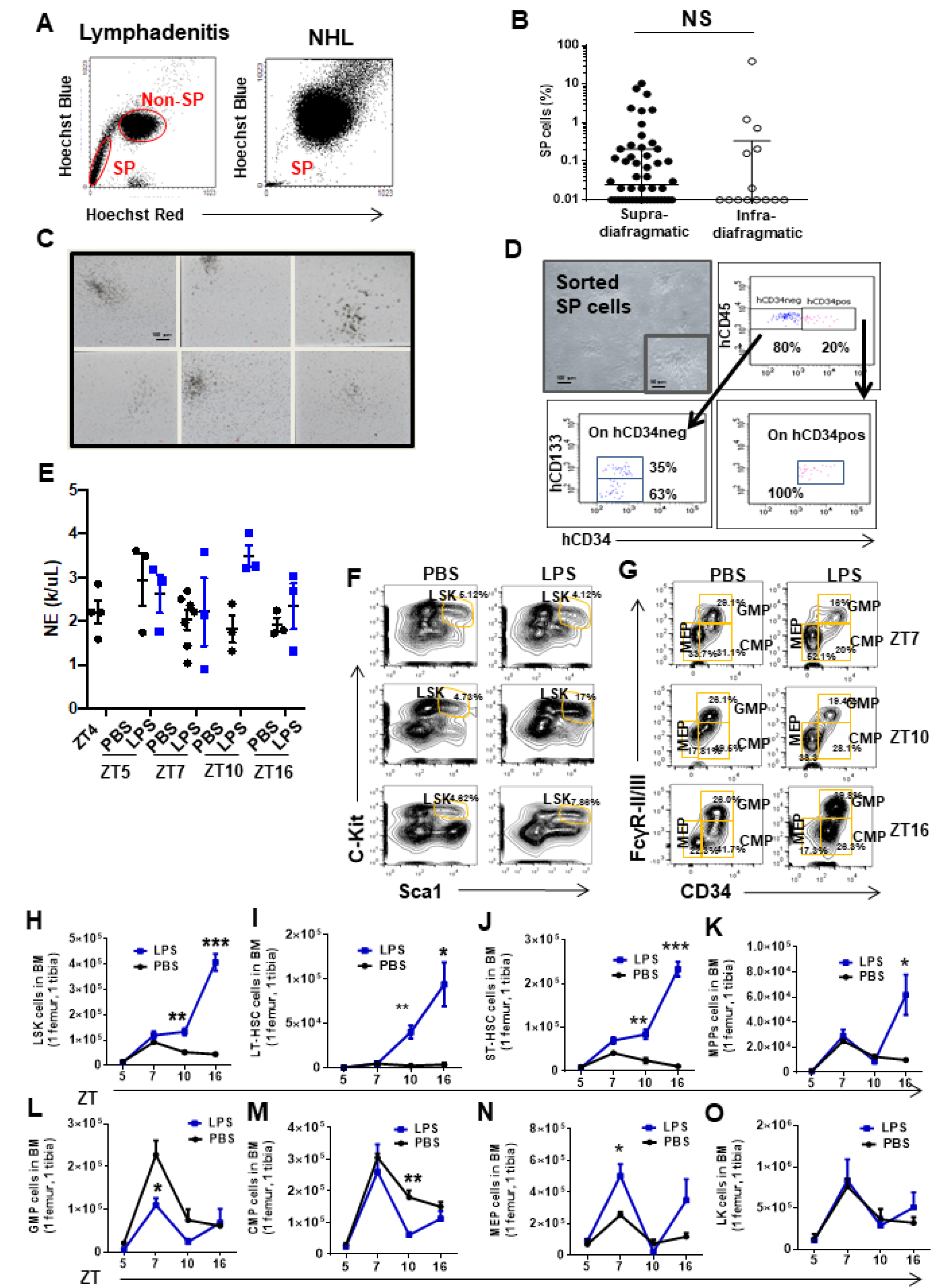
Clonogenic and long-term multilineage potential of human and murine HSC/P in LN. (A) Representative flow cytometry dot plots of SP cells from human LN biopsies diagnosed with lymphadenitis (left flow panel) and non-Hodgkin Lymphoma (NHL, right flow panel). SP cells form a tail cluster on the left side. (B) Graph represents the percentage of SP cells in human LN biopsies categorized according to their anatomical location. Supra-diafragmatic location (solid circles) included neck and axillary LN and infra-diafragmatic location (open circles) included mesenteric and inguinal LN. (C) Representative colony-forming units (CFU-C) micrographs in human LN diagnosed with follicular lymphadenitis. Scale bar: 100 µm. (D) Clonogenic potential of lymphadenitis-derived SP cells (upper left; with magnification of a CFU-GM in outlined inset) and FACS analysis of LN SP-derived hematopoietic progenitors (upper right and bottom panels) maintained for 1 week in culture as previously described(*19*). All SP-derived progenitors were positive for pan-leukocyte surface marker CD45 but had heterogenic expression for CD34 and CD133 surface markers. (E) Absolute neutrophil count in PB from C57Bl/6 mice pre-treated with PBS (black circles) or LPS (blue circles) at different circadian cycle times. (F-G). Representative FACS profiles of murine BM HSC/P in response to *in vivo* administration of PBS (left panels) or LPS (right panels) at specified ZT. (H-O) Time response in the BM content of LSK (H), LT-HSC (I), ST-HSC (J), MPP (K), GMPs (L), CMP (\M), MEP (N) and LK cells (O) in response to PBS (black lines) or LPS (blue lines). Values represent mean ± SD of a minimum of 4 mice per group. *P<0.05, **P<0.01, ***P<0.001.

**Figure S2.**
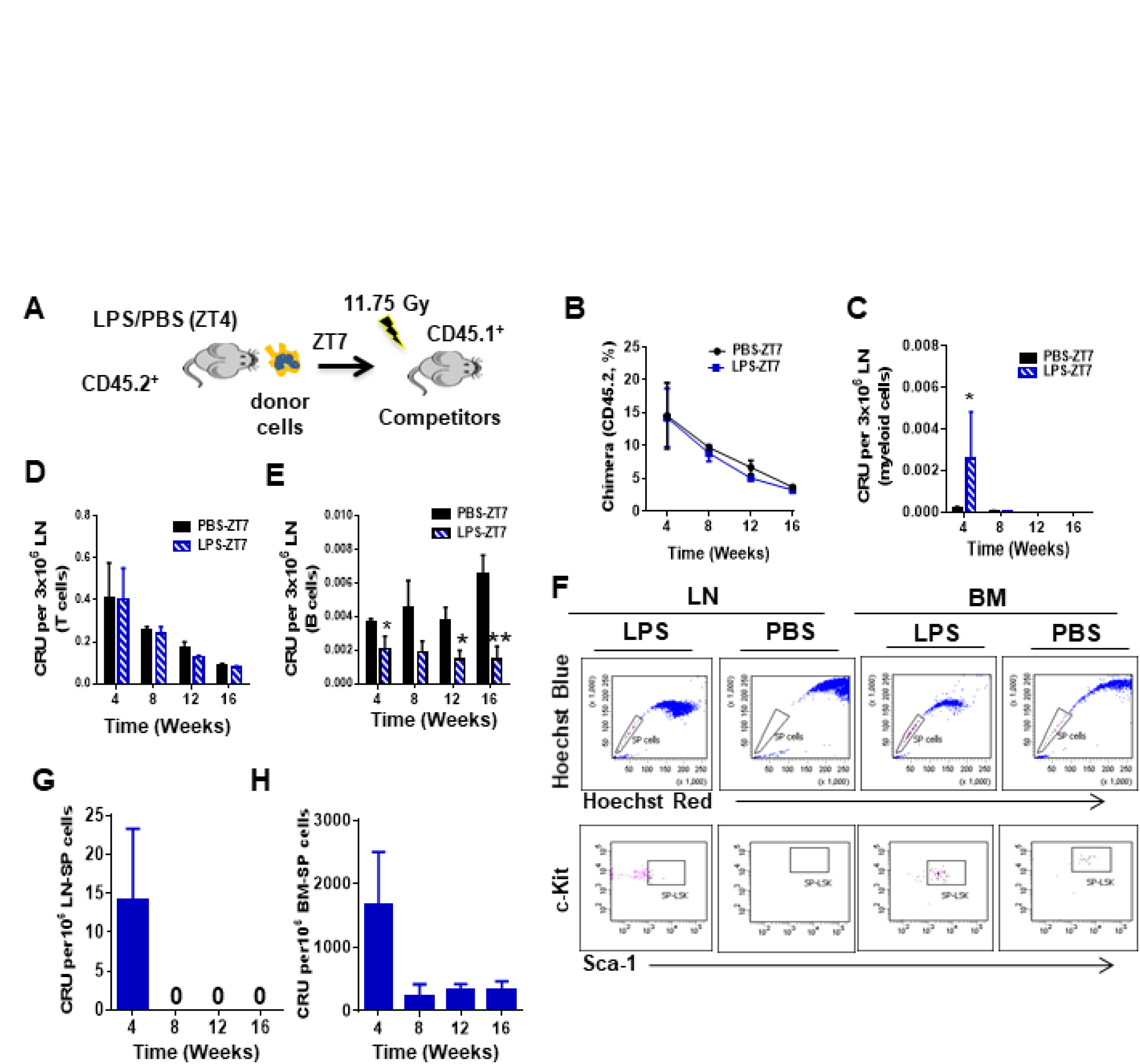
Myeloid progenitor migration to LN in response to LPS is independent of NFκB activation. (A) Schema for the competitive long-term reconstitution potential of LN cells (donor cells) from C57Bl/6 (CD45.2^+^) mice treated with PBS/LPS and harvested at ZT7(3h), into lethally irradiated CD45.1^+^ B6.SJL^Ptprca Pep3b/BoyJ^ recipient mice (n=3 mice per group). (B-E) Competitive repopulating unit (CRU) assay of LN cells as assessed by flow cytometry of allotype CD45.2 expressing cells in transplanted mice followed for up to 16 weeks. (B) Overall CRU as gated on CD45.2^+^ cells in response to donor cells from mice pre-treated with PBS (black line) or LPS (blue line).(C-E) Myeloid cell (C), T-cell (D) and B-cell (E) contributing CRU in response to donor cells from mice pre-treated with PBS (solid bar) or LPS (blue mosaic bar). (F) Representative example of FACS analysis for SP cells (upper dot plots) and SP-LSK potential (lower dot plots) from PBS- and LPS-treated LN or BM tissues at ZT7 (3h). Hoechst: Hoechst 33342 staining for blue (blue) and red (red) fluorescence emissions. (G-H) Donor-derived chimera of sorted LN-or BM-SP cells obtained from LPS-treated mice.

**Figure S3.**
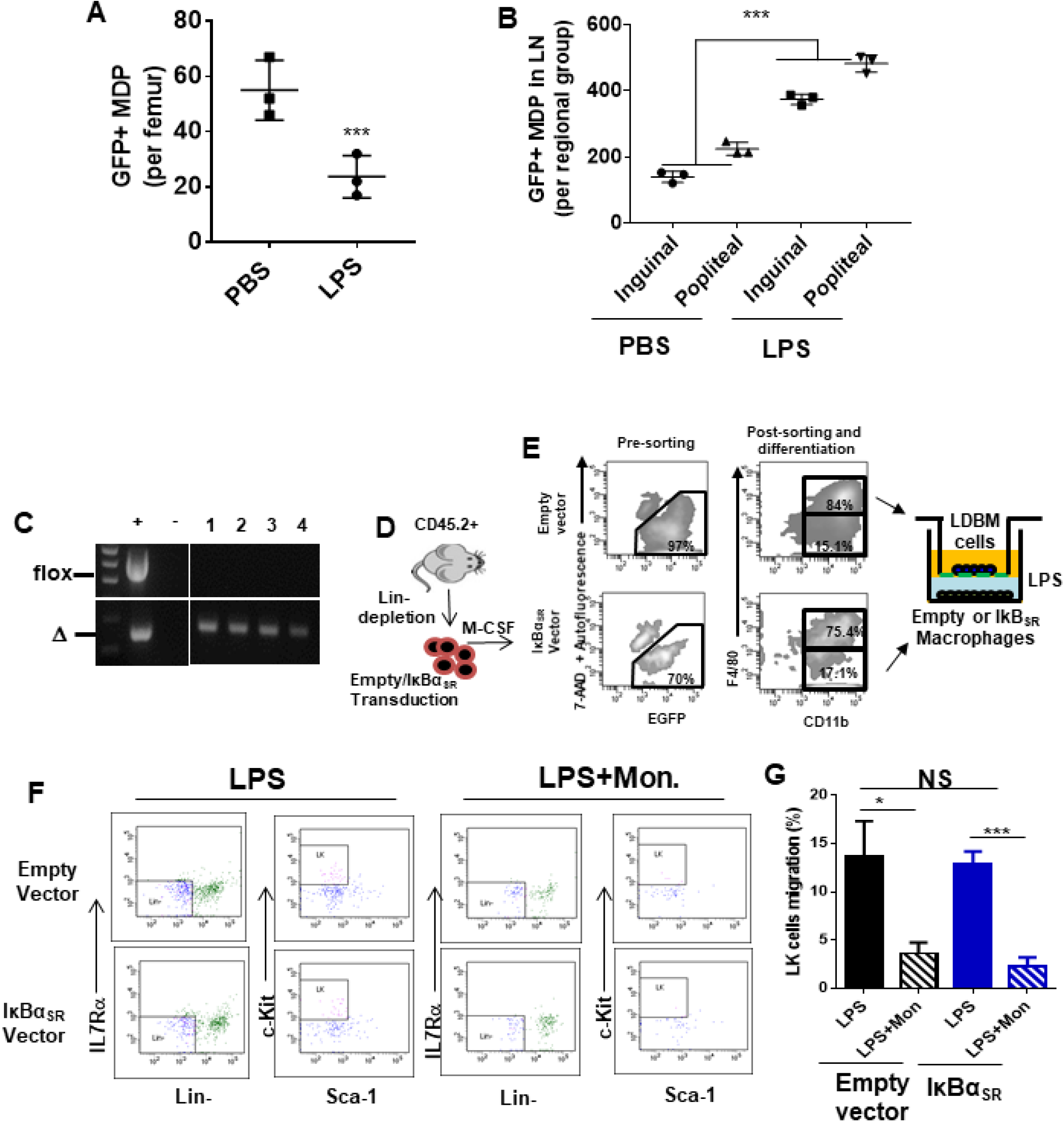
Myeloid progenitor migration to LN in response to LPS is independent of NFκB activation. (A) Flow cytometry analysis of GFP MDPs in bone marrow showing decreased MDPs upon LPS treatment. (B) Flow cytometry analysis of migrated GFP MDPs in lymph node (per regional group) showing increased GFP^+^MDPs in Inguinal and Popliteal lymph nodes. (Black bar EGFP^+^ and Blue Bar EGFP^−^. (C) Representative PCR amplifications show Traf6 deleted in circulating cells in four *Mx1cre;Traf6^flox/flox^* mice after poly(I:C) injections (10mg/Kg/2 days x 5 doses). (D) Schema for BM-derived macrophages and exogenous expression of IκBα mutant which is resistant to proteasome degradation. BM Lin^−^ cells were transduced with Pmscv-puro-eGFP bicistronic retroviral vector encoding full length of IκBα mutant and GFP^+^ protein. EGFP^+^ Lin^−^ cells were sorted and differentiated in culture by M-CSF cytokine to macrophages. Green fluorescent CD11b+-macrophages were layered on bottom chamber and stimulated them with LPS to generate myeloid chemotaxis gradient. (E) FACS strategy for migrated LK (Lin^−^/c- Kit^+^/Sca1^−^) cells contained in LDBM from upper chamber to the transduced macrophage bottom stimulated with LPS and LPS+monensin (LPS+Mon) for 4 hours as depicted in J. (F) Representative FACS dot plots demonstrating gating strategy to identify migrating GMP populations from the transwell migration assays. (G) Graph represents LK cell migration towards transduced macrophages in presence of LPS (solid bars) or LPS+Mon (mosaic bars). Values represent mean±SD of three mice and per triplicate. NS: not significant. *P<0.05 ***P<0.001.

**Figure S4.**
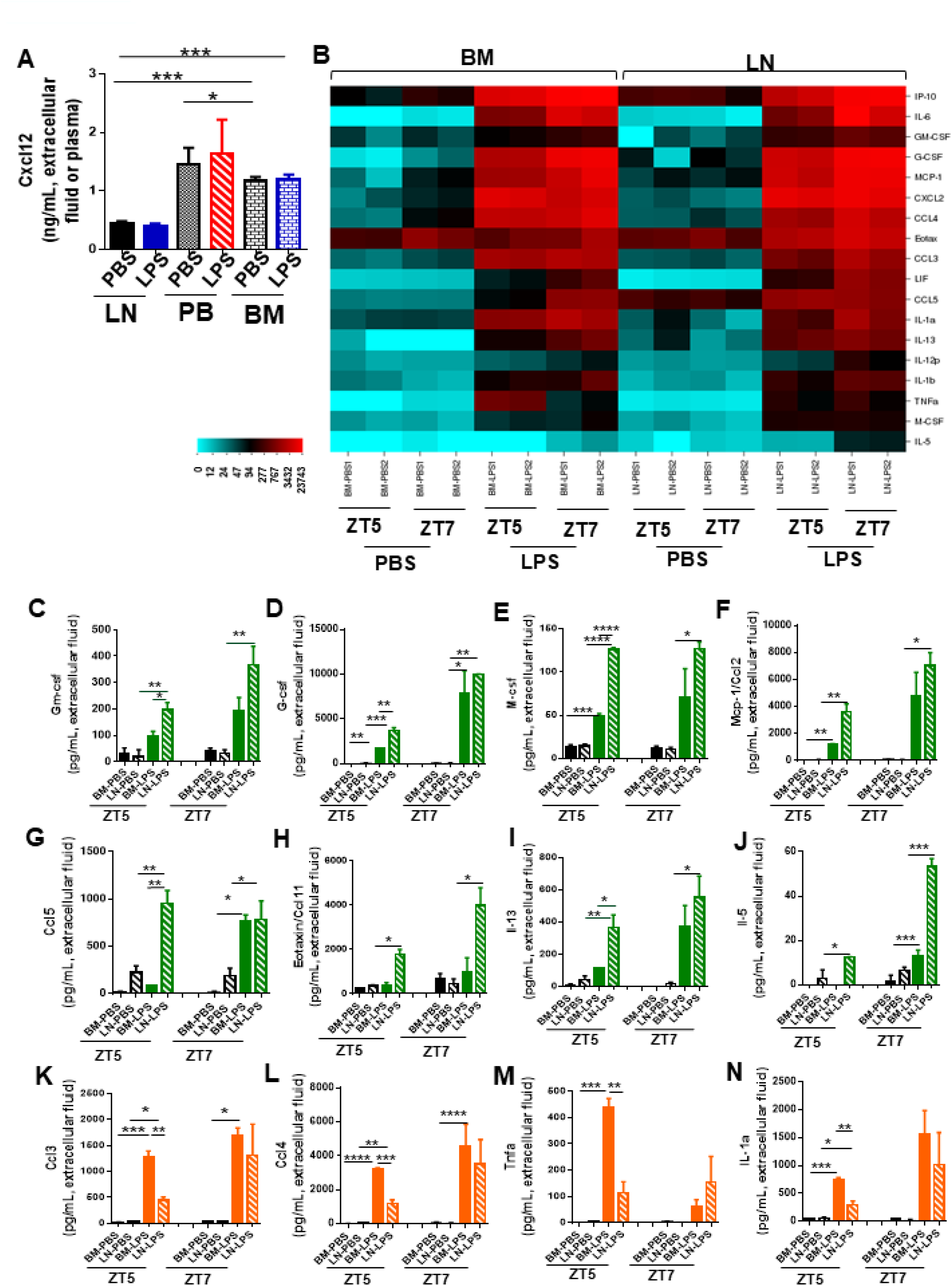
Inflammation induces temporal changes in chemokine and cytokine signatures in BM and LN. (A) Cxcl12 in femoral or LN extracellular fluid and plasma after PBS/LPS *in vivo* administration at ZT7 (3h). (B) Heat map showing cytokine profiling release into the extracellular fluid of femora and LN in response to PBS/LPS at ZT5 (1h) and ZT7 (3h). (C-O). Graphs represent levels of relevant cytokines and chemokines associated with migration/inflammatory response and released into LN extracellular fluid (black and green bars) or into femoral extracellular fluid (black and orange bars) after PBS/LPS administration into C57Bl/6 mice at ZT5 (1h) and ZT7 (3h). (C-J) Extracellular LN levels of Gm-csf (C), G-csf (D), M-Csf (E), Mcp-1 (F), Ccl5 (G), Eotaxin/Ccl11 (H), IL-13 (I), IL-5 (J). (K-N) Extracellular BM levels of Ccl3 (K), Ccl4 (L), Tnfα (M) and IL-1α (N). Values are mean ± SE of two mice per treatment and experiment, pooled from two independent experiments *P<0.05, **P<0.01, ***P<0.001, ****P<0.0001.

**Figure S5.**
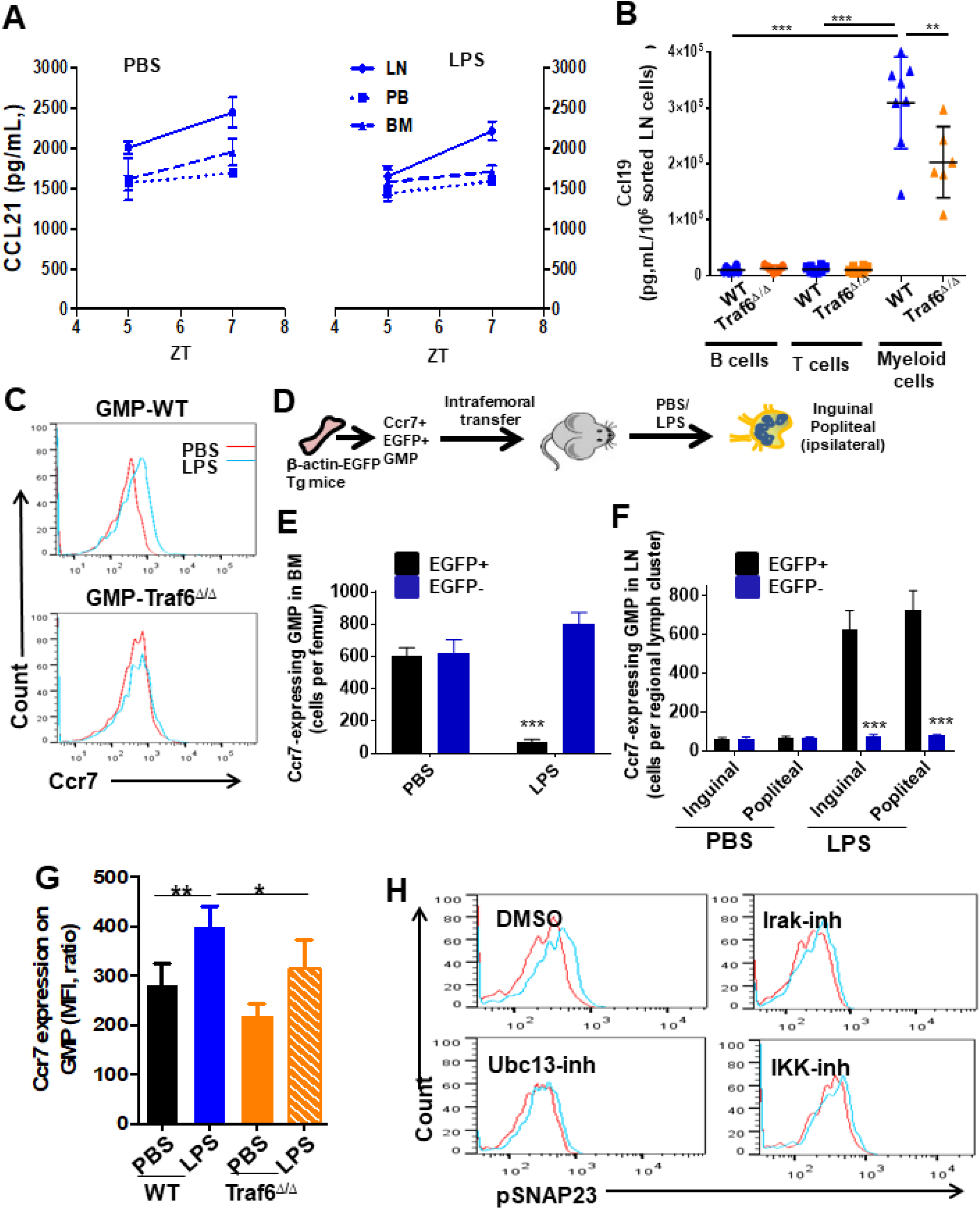
Myeloid expression of Ccl19 ligand in LN and short-term differentiation of pre-treated GMPs. (A) Ccl21 in femoral or LN extracellular fluid and PB plasma after PBS (left lines) or LPS (right lines) administration at different circadian cycle times. (B) Ccl19 released into the supernatant from sorted WT and Traf6-deficient LN T^+^ cells, B^+^ cells and CD11b^+^ myeloid cells after LPS stimulation *in vitro*. Values represent mean ± SEM of two independent experiments. **P<0.01, ***P<0.001. (C-D). Membrane Ccr7 expression on GMP at ZT5 (1h) after PBS/LPS administration into WT and Traf6 deficient mice shows significant differences between groups. (C) Representative overlap histograms of Ccr7 expression on GMP-WT cells (upper) and GMP-Traf6^Δ/Δ^ (lower) cells after PBS (red line) or LPS (blue line). (D) Schema of transfer of Ccr7+ EGFP+ BM cells from b-actin-EGFP+ transgenic animals to femurs of WT recipient mice and administered PBS or LPS. (E-F) Femoral content of Ccr7-expressing GMP cells in BM (E) and regional (inguinal and popliteal) LN (F). (G) MFI of Ccr7 on GMPs from WT and Traf6 deficient mice in presence of PBS (black and orange solid bars) or LPS (blue solid bar and orange mosaic bar). Values represent mean mean ± SD of three mice. (H) Representative overlap histograms of phospho-SNAP23 (pSNAP23) into LN myeloid cells treated with DMSO (upper left), Irak1/4-inh (upper right), Ubc13-inh (lower left) and IKK-inh (lower right) and stimulated with PBS (red lines) or LPS (blue lines).

**Figure S6.**
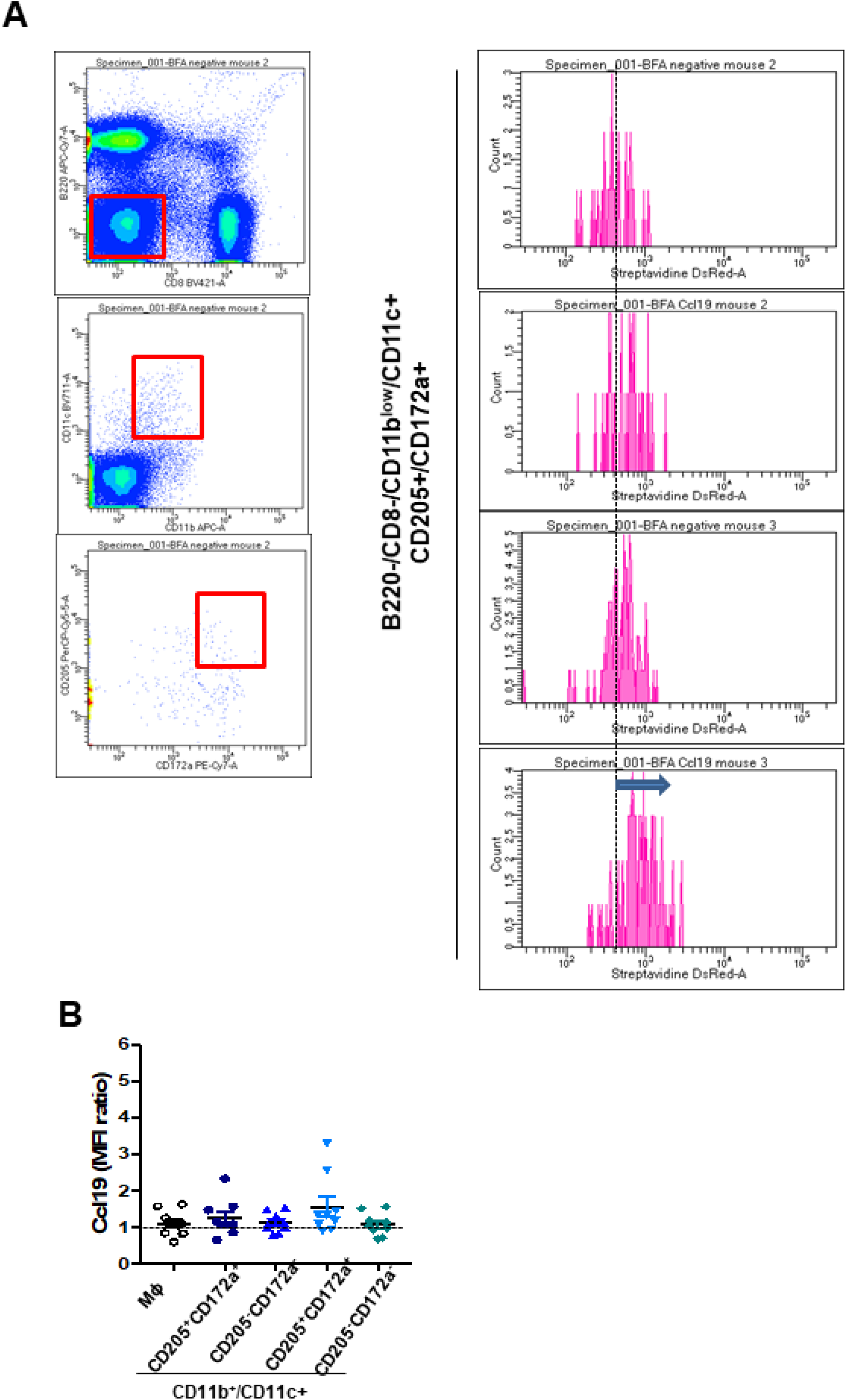
Basal Ccl19 detection into LN populations. (A) Gating strategy to determine Ccl19 pre-stored into LN-residing myeloid cells basally. (B-C) Mean fluorescence intensity (MFI) quantification of pre-stored Ccl19 into LN-residing CD11b^low^/CD11c^+^ cDC (B) and pDC (C) populations in basal conditions.

## Supplementary Movies

**Movie S1 caption**

Representative example of multi-photon microscopy processed with Imaris software of a femur from one Lyve1-EGFP mouse treated with PBS at 1 hour after administration (ZT5). Bone tissue is identified as second-harmonic (SHG) signal (blue). Imaris software was used to measure distance between DiI labeled cells and GFP positive lymphatic vessels using 3D images.

**Movie S2 caption**

Representative example of multi-photon microscopy processed with Imaris software of a femur from one Lyve1-EGFP mouse treated with LPS at 1 hour after administration (ZT5). Bone tissue is identified as second-harmonic (SHG) signal (blue). Imaris software was used to measure distance between DiI labeled cells and GFP positive lymphatic vessels using 3D images.

**Movie S3 caption**

Representative example of transcortical lymphatics discovered by staining with anti-Lyve1 (red) in wild-type C57Bl/6 mice. Bone tissue is identified as second-harmonic (SHG) signal (blue). Green signal is autofluorescence.

